# Predicting glycan structure from tandem mass spectrometry via deep learning

**DOI:** 10.1101/2023.06.13.544793

**Authors:** James Urban, Chunsheng Jin, Kristina A. Thomsson, Niclas G. Karlsson, Callum M. Ives, Elisa Fadda, Daniel Bojar

## Abstract

Glycans constitute the most complicated post-translational modification, modulating protein activity in health and disease. However, structural annotation from tandem mass spectrometry data is a bottleneck in glycomics, preventing high-throughput endeavors and relegating glycomics to a few experts. Trained on a newly curated set of 300,000 annotated MS/MS spectra, we present CandyCrunch, a dilated residual neural network predicting glycan structure from raw LC-MS/MS data in seconds (Top1 Accuracy: 87.7%). We developed an open-access Python-based workflow of raw data conversion and prediction, followed by automated curation and fragment annotation, with predictions recapitulating and extending expert annotation. We demonstrate that this can be used for *de novo* annotation, diagnostic fragment identification, and high-throughput glycomics. For maximum impact, this entire pipeline is tightly interlaced with our glycowork platform and can be easily tested at https://colab.research.google.com/github/BojarLab/CandyCrunch/blob/main/CandyCru nch.ipynb. We envision CandyCrunch to democratize structural glycomics and the elucidation of biological roles of glycans.

## Main

As the most abundant post-translational modification, glycans are frequently dysregulated and mechanistically involved in diseases ranging from cancer^1^ to metabolic disorders^2^. The exact structure of complex carbohydrates is often key in mediating their function^3^, such as sialic acid only facilitating influenza infection in a particular linkage orientation^4^. From biomarkers to mechanistic understanding^1,2^, structural resolution thus is relevant for integrating and using glycan information for biomedical gains. In the context of systems biology, glycans are routinely measured via mass spectrometry-based glycomics^5^, providing insights into which structures or substructures are dysregulated, which can be further analyzed with various methods^6,7^.

Currently, structural determination of glycans is, at best, semi-manual and proceeds structure by structure^8^. Since different glycan structures can result in the same mass, structural isomers are routinely separated via liquid chromatography (LC)^9^, followed by fragmentation into smaller sub-structures by MS, conceptually akin to shotgun sequencing. Current in-depth workflows are hard to parallelize, with a general trade-off between resolution and scale^10^. All this has relegated structural glycomics to a few experts, inaccessible to most life science researchers.

Extensive work by Harvey and others^8,11,12^ has demonstrated that, in principle, most substructures^13^, linkages^14^, and monosaccharides^15^ have diagnostic fragments or intensity ratios. Using this fine structural information that is contained within MS/MS spectra, along with basic biosynthetic assumptions, it is thus frequently possible to achieve high resolution annotations of native glycans. In practice, however, annotation is often restricted to essentially topological assignments, not least due to time-constraints. Nuances of diagnostic indicators are challenging for humans to decrypt manually or encode programmatically, especially at scale and accommodating diverse experimental setups, as each linkage and monosaccharide can be affected by its sequence context^16^. This combinatorial explosion, combined with rich data, is promising for scalable artificial intelligence approaches, which can learn complex mapping functions, as recently demonstrated by endeavors such as AlphaFold2^17^.

So far, computational attempts to automate MS based glycomics^18–23^ did not engage with deep learning. Rather, they relied on various search methods, to either search for possible topologies given a precursor ion mass or suitable reference spectra, loose constraints that may yield unphysiological predictions. Their primary limitations are scale and annotation resolution, ranging from composition to glycan topology. Neither linkage type nor monosaccharide stereoisomers are commonly resolved during this algorithmic sequencing. Additional hurdles to their wider adoption include poor generalizability, as none of them employ a rigorous train-test mentality, a standard practice in machine learning to evaluate methods on held-out data to prevent overfitting. Many tools were designed for very specific problems and were often tested on few spectra^18–20^, precluding their usage in many experimental setups.

Recent efforts in related fields, particularly in proteomics^24,25^, have employed scalable deep learning strategies in mass spectrometry analysis. Proteomics has partially similar challenges to glycomics, e.g., precursor structure elucidation given fragment ions. We thus posit that the translation of analogous methods to structural glycomics, combined with domain-inspired additions such as biosynthetic constraints and building on the accumulated work of many years of glycomics analysts, could be a major leap forward for the field and the usage of glycomics in the broader life sciences.

We present a scalable and accurate workflow for predicting glycan structure from LC-MS/MS data, centered on our deep learning model CandyCrunch. Using a large-scale, curated set of tandem spectra from diverse experimental set-ups, CandyCrunch predicts glycan structure with high accuracy (>87%), outperforms existing methods on this task, and matches/extends expert annotations on unseen data. This is facilitated by various domain-specific advancements, e.g., considering glycan structure similarity in the loss function. We embedded this into a downstream workflow converting predictions into interpretable results, further reducing false-positive rates, and estimating relative abundances; all in seconds. This workflow includes CandyCrumbs, a comprehensive MS/MS fragment annotation plug-in we developed here. We used this to uncover diagnostic fragments and more complex fragmentation behavior at scale, underpinned by molecular dynamics simulations. Finally, we annotate novel glycomes, analyze biosynthetic constraints at scale, and demonstrate that our pipeline can be used in high-throughput glycomics. Our methods are accessible within a Python package (https://github.com/BojarLab/CandyCrunch) and a free-standing Google Colab notebook at https://colab.research.google.com/github/BojarLab/CandyCrunch/blob/main/CandyCrunch.ip ynb.

## Results

### CandyCrunch predicts glycan structure within a domain knowledge-suffused workflow

Reasoning that the fragmentation patterns and propensities (i.e., intensity ratios) in tandem mass spectrometry are predictive of glycan structure – a relationship that is used by human experts in annotation – we set out to learn this association via machine learning. For this, we collected and curated an unprecedentedly large set of annotated LC-MS/MS spectra that derive from glycans (Fig. 1A-B; see Methods). We envision that, even beyond our efforts here, this dataset will be a valuable resource for data-driven approaches in glycomics. Crucially, this dataset aims to provide a representative view over current glycomics data, with a total of nearly 500,000 labeled MS/MS spectra from >2,000 glycomics experiments, encompassing all major eukaryotic glycan classes (*N*-linked, *O*-linked, glycosphingolipid, milk oligosaccharides) and the most common experimental setups for glycomics. The exact composition of this dataset, broken down by glycan classes and experimental parameters can be found in Supplementary Table 1. To avoid overrepresenting some classes (e.g., core 1 *O*-glycan), we then limited each class to a maximum of 1,000 example spectra (see Methods for details) and used the resulting 300,000 spectra to train our model the most likely glycans in a multi-class classification set-up (see Methods).

**Figure 1.**
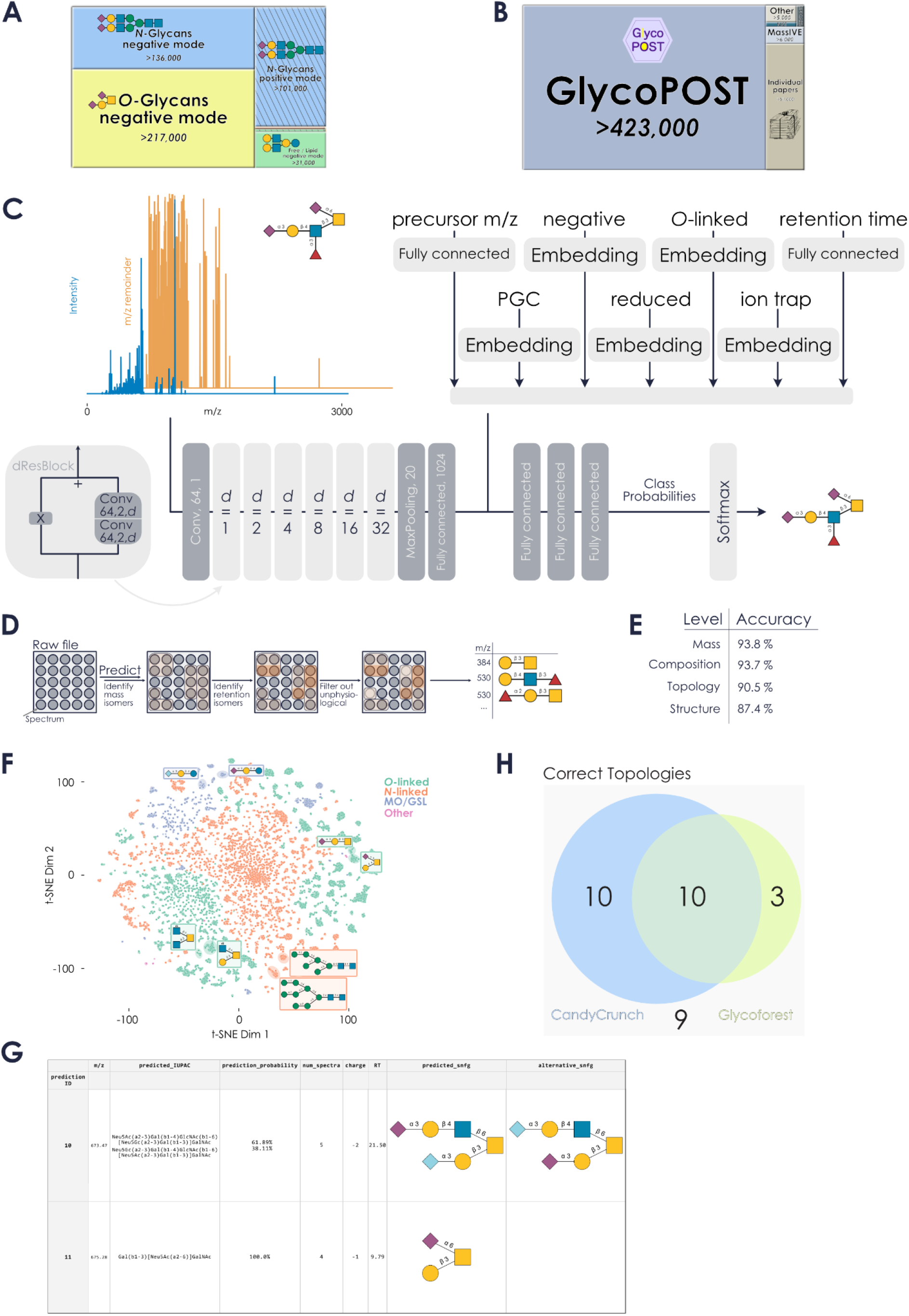
Predicting glycan structure via deep learning. **A-B)** Overview of the curated dataset of glycomics LC-MS/MS by glycan class (A) and source (B). Diagonal bars indicate positive ion mode data. The numbers correspond to spectra with annotations. **C)** Schematic view of CandyCrunch model architecture. **D)** Pipeline of curating glycan predictions from raw file to final output table. **E)** Evaluating top-1 accuracy on the independent test-set (see Methods) across different levels of resolution. **F)** Learned representations of all spectra in the test-set are shown via t-SNE, colored by glycan class. Examples are illustrated with their glycan structure. **G)** Excerpt from an example prediction output using our Colab notebook on the file JC_171002Y1.mzML^26^. **H)** Proportional Venn diagram of the comparison of CandyCrunch and Glycoforest on the raw file JC_131210PMpx5.mzML^18^, not used for training CandyCrunch but used for developing Glycoforest. Shown are topologies (Glycoforest does not output full structures) matching those of the human annotator for each model (see Supplementary Fig. 5 for detailed comparison). All shown masses are from reduced glycans. Glycans here and in the entire manuscript are drawn using GlycoDraw^27^ according to the Symbol Nomenclature for Glycans (SNFG).

This resulted in our dilated residual neural network CandyCrunch, a model architecture suited to mass spectrometry data^25^. Since experimental parameters such as the ion mode drastically change fragmentation patterns, it uses the MS/MS spectrum, retention time, precursor ion m/z, and experimental parameters (e.g., LC type, ion mode, etc.) as input and predicts glycan rankings as its output (Fig. 1C). We note that we neither claim, nor sought to obtain, the most frugal model for this task, but rather the most performant and flexible, without noticeable hardware limitations (CandyCrunch can be readily used on a typical laptop). The model is part of a pipeline applied to a raw file (e.g., .mzML or .mzXML files), which groups predictions based on mass-and retention isomers and further curates predictions with, e.g., diagnostic ions (Fig. 1D).

If precursor ion intensities are available in the raw file, this pipeline can also estimate relative abundances. These abundances correlate well with those gained by LC peak area integration (Supplementary Fig. 1), a state-of-the-art approach for estimating relative abundances. Overall, CandyCrunch is highly performant, with an accuracy of >87% of the top-ranked structure prediction in the independent test set (Fig. 1E), performing comparably across glycans (Supplementary Fig. 2A-B) and across different MS setups and glycan classes (Supplementary Table 2). Custom loss functions estimating structural distance to the ground truth, and many more domain-knowledge-inspired modifications (see Methods), ensure that even erroneous predictions are structurally close to the correct solution. Our approach also includes incompletely resolved structures, so that prediction uncertainty can be meaningfully conveyed via, for instance, missing linkage information (indicated by a higher topology accuracy than structure accuracy; Fig. 1E).

Learned representations of spectra by CandyCrunch cluster by glycan sequence and glycan class (Fig. 1F), demonstrating that the model has learned to accommodate experimental variability. Further, structurally related glycans, even within the same class, tend to cluster together in the learned representation space. This can be quantified by comparing the cosine distance of learned representations of pairs of glycans with their structural distance, revealing that the co-clustering described by the representations is indeed suggestive of structural relatedness of glycans (two-sided Mantel test of correlating the two resulting cosine distance matrices; p < 0.001).

In framing CandyCrunch as a multiclass classification problem (i.e., ranking the likelihood of pre-defined glycans), we minimized the chance for unphysiological glycans in the output, a very real possibility otherwise, given the sparsity of real glycan sequences among possible sequences^28^. However, this made zero-shot predictions – predicting a glycan sequence that was absent from our training set – conceptually infeasible. As repositories such as GlycoPOST do not catalogue all physiological glycans, and glycomics studies, such as mucin-type *O*-glycomics^29^ or milk glycomics^30^, routinely discover new structures, we set out to augment our pipeline to allow for, limited, zero-shot prediction outside our 3,508 defined glycans.

Reasoning that glycans in a biological sample tend to be biosynthetically related, i.e., contain precursors/intermediates of larger biosynthetic pathways, we turned to our recently developed method of constructing glycan biosynthetic networks^7^. Applying this method to a typical CandyCrunch output (Supplementary Fig. 3) revealed the existence of necessary intermediate structures that were absent from our predictions but would explain spectra without a valid prediction. We thus added this routine as an optional step in our inference workflow, to facilitate a certain subset of physiological zero-shot predictions.

CandyCrunch is fundamentally database-independent but can be further enhanced by methods leveraging databases, such as defined within glycowork^31^, to augment predictions downstream. By carefully selecting a suitable subset of reference structures (e.g., by taxonomy, glycan class, or tissue), matches for unexplainable spectra could be proposed. These potential matches were then cross-checked for diagnostic ions as well as ranked by biosynthetic compatibility with true predictions. This, again, allowed for a certain subset of zero-shot predictions. It should be noted that this procedure still balanced the theoretical constraint of physiological glycans with the reality of encountering novel structures in biological samples. Our final inference workflow then also contained this latter expansion, resulting in a ranked prediction output that can be further investigated by the researcher (Fig. 1G).

Next, we compared CandyCrunch with alternative approaches to this problem. As a preface, we should note that no current approach combines CandyCrunch’s advantages of scale, generalizability, performance, and its flexibility in usage (Supplementary Fig. 4A). Further, most methods are only maintained for the briefest of periods and are no longer realistically accessible. Thus, we had to effectively constrain ourselves to compare CandyCrunch on individual raw files that were specifically used to build these alternative approaches, while we excluded them during training. Still, in direct comparison with state-of-the-art methods such as Glycoforest^18^ on challenging fish mucin glycans, CandyCrunch demonstrated a greater overlap with manual expert annotations (Fig. 1H; 62.5% versus Glycoforest’s 40.6%) and a substantially higher structural resolution (Supplementary Fig. 5). In addition, by tethering CandyCrunch and the below-mentioned CandyCrumbs to our glycowork ecosystem^31^ and by providing everything open-source, we substantially increase the chances for the long-term viability of our presented methods.

Applied to fully unseen datasets, CandyCrunch routinely achieved high performance (Supplementary Table 3; topology: 92% accuracy, structure: 84% accuracy) and potentially can extend expert annotations by correctly capturing additional structures and isomers (Supplementary Fig. 6). The additional predictions in this sample partly even stemmed from remnant glycans from the previous sample, showcasing the exceptional sensitivity of our model. We would like to highlight here that the cross-training of CandyCrunch on all glycan classes yielded performance synergy, as a model only trained on *O*-glycans performed worse for predicting *O*-glycans (Supplementary Table 3; topology: 84% accuracy, structure: 79% accuracy) than the model trained on all classes. We posit that this was due to the structure-based loss function we used for training, as well as shared information between spectra of different classes, stemming from shared glycan motifs across classes (e.g., Neu5Ac-Hex).

The speed and relatively low resource requirements of CandyCrunch (Supplementary Fig. 7A) mean that samples can be exhaustively analyzed, without practical constraints to the X most abundant structures, which is a routine necessity in human analysis. In its typical application, CandyCrunch also makes fewer assumptions about what is or should be present in a sample, enhancing the chances for novel discoveries. This means, e.g., that co-released *N*-glycans can be detected in *O*-glycan preparations (Supplementary Fig. 8).

### CandyCrumbs automatically annotates fragment ions and facilitates diagnostic ion discovery

When analyzed by humans, fragment ions are usually annotated via the Domon-Costello nomenclature^32^ and used for elucidating the structure of a glycan. While there are programs that automate this assignment^21,22^, they are either only accessible via graphical user interfaces or only provide annotations for simple fragment ions. We thus decided to implement an exhaustive Python-based solution to this problem, CandyCrumbs, which is also freely available via the CandyCrunch Python package. Given a candidate glycan sequence and fragment peaks, CandyCrumbs can automatically and rapidly (Supplementary Fig. 7B) annotate fragment ions in Domon-Costello and IUPAC-condensed nomenclature (Fig. 2A, see Methods). Compared to alternative approaches, this presents the most feature-complete and rapid implementation of this task (Supplementary Fig. 4B).

**Figure 2.**
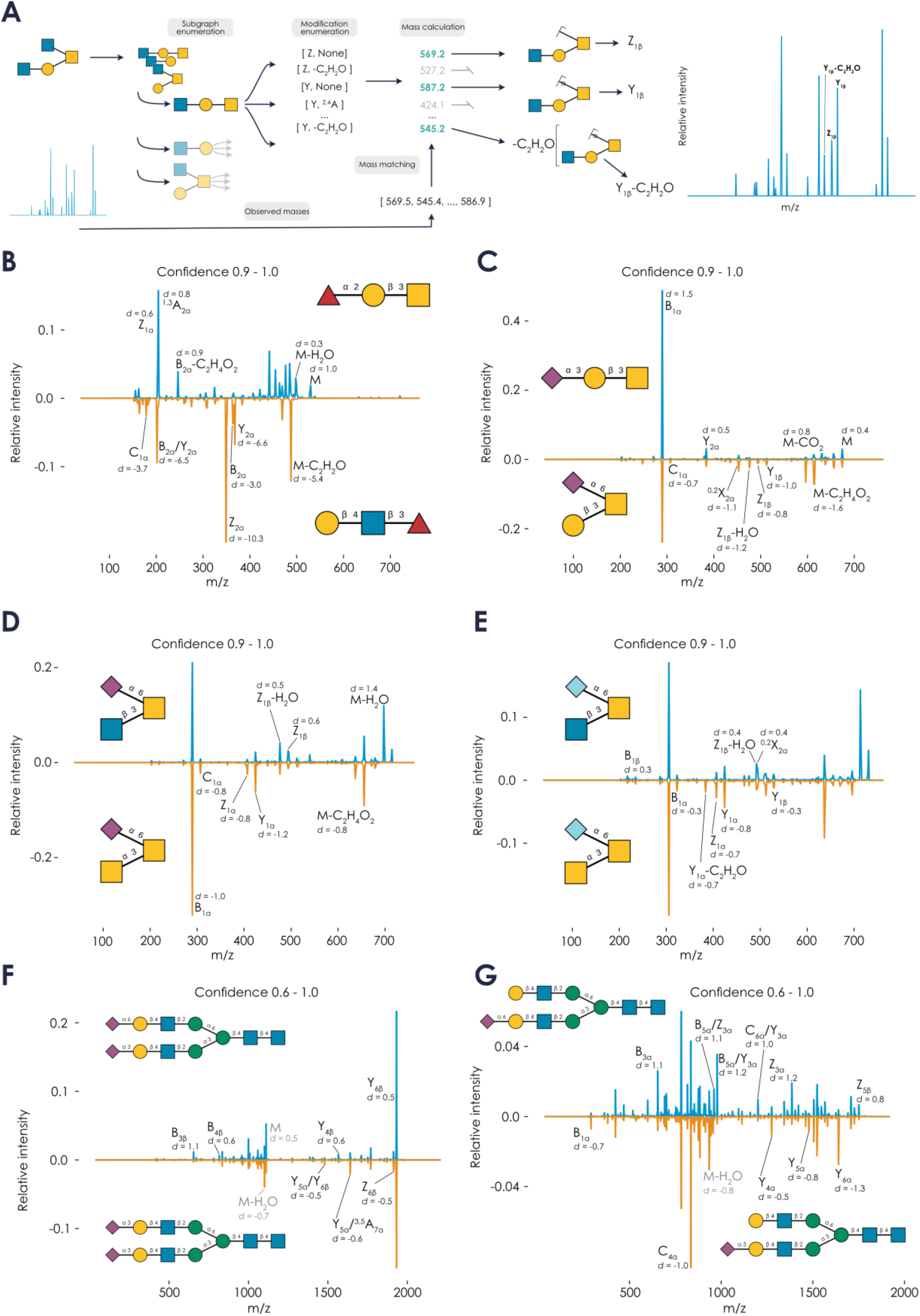
Discovering diagnostic fragmentation using CandyCrumbs. **A)** Schematic view of the CandyCrumbs workflow for automatic fragment ion annotation. **B-E)** Negative ion mode spectra of reduced glycans with prediction confidence between 0.9 and 1.0 for Fucα1-2Galβ1-3GalNAc / Galβ1-4GlcNAcβ1-3Fuc (A), Neu5Acα2-3Galβ1-3GalNAc / Galβ1-3(Neu5Acα2-6)GalNAc (B), GlcNAcβ1-3(Neu5Acα2-6)GalNAc / GalNAcα1-3(Neu5Acα2-6)GalNAc (C), and GlcNAcβ1-3(Neu5Gcα2-6)GalNAc / GalNAcα1-3(Neu5Gcα2-6)GalNAc (D) were averaged and juxtaposed. Fragments exhibiting differential abundance were labeled by CandyCrumbs in the Domon-Costello nomenclature^32^. **F-G)** Negative ion mode spectra of reduced glycans with prediction confidence between 0.6 and 1.0 for Neu5Acα2-3Galβ1-4GlcNAcβ1-2Manα1-3(Neu5Acα2-6Galβ1-4GlcNAcβ1-2Manα1-6)Manβ1-4GlcNAcβ1-4GlcNAc / Neu5Acα2-3Galβ1-4GlcNAcβ1-2Manα1-3(Neu5Acα2-6Galβ1-4GlcNAcβ1-2Manα1-6)Manβ1-4GlcNAcβ1-4GlcNAc (F) and Neu5Acα2-6Galβ1-4GlcNAcβ1-2Manα1-3(Galβ1-4GlcNAcβ1-2Manα1-6)Manβ1-4GlcNAcβ1-4GlcNAc / Neu5Acα2-3Galβ1-4GlcNAcβ1-2Manα1-3(Galβ1-4GlcNAcβ1-2Manα1-6)Manβ1-4GlcNAcβ1-4GlcNAc (G) were averaged, juxtaposed, and labeled similar to (B-E). Doubly-charged fragment ions are colored gray.

Further, we used several domain knowledge-inspired heuristics and probability rules to highlight the most probable fragments (Supplementary Fig. 9; see Methods), if multiple fragmentation options could result in an m/z value that is acceptable at a given threshold. We then also integrated CandyCrumbs within the aforementioned open-access Colab notebook (at https://colab.research.google.com/github/BojarLab/CandyCrunch/blob/main/CandyCrunch.ip ynb) for full flexibility. Our implementation of CandyCrumbs then allowed us to use it in a high-throughput setting and integrate it into CandyCrunch workflows, such as for identifying diagnostic ions at scale as discussed below, to aid expert annotation of challenging cases.

Reference spectra are routinely used as high-quality examples in semi-manual annotation^33^. As “spectrum quality” is an ill-defined and subjective characteristic, we aimed to quantify this aspect by using calibrated^34^ prediction confidence of CandyCrunch as a proxy, with the reasoning that a more confidently assessed spectrum is a higher-quality spectrum with more information for effective prediction. Rather than one reference spectrum, i.e., the usual approach^33^, we then extracted hundreds to thousands of high-quality spectra for a given structure from our dataset and engaged in highly-powered statistical comparisons between isomers. This identified numerous diagnostic ions and/or ratios for topologically distinct (Fig. 2B-C) and identical (Fig. 2D-E) isomers, with large effect sizes. This also extended to other glycan classes and, e.g., facilitated detecting conserved fragmentation differences of linkages (e.g., stronger B3 ion in α2-6 vs α2-3) across glycan backbones (Fig. 2F-G) and recapitulated known effects from the literature^35^, such as a higher stability of α2-6 vs α2-3 in negative mode (see B1 ion in Fig. 2G). Importantly, these differences diminished, and eventually vanished, with lower-quality spectra (Supplementary Fig. 10). We then analyzed the predictiveness of these diagnostic features when reducing spectrum quality. Intriguingly, some diagnostic features, even if they were not the strongest initial signal, remained predictive even for medium-to low-quality spectra (Supplementary Fig. 11), making them promising candidates for aiding annotation.

Similarities between Neu5Ac- and Neu5Gc-versions of the same isomers (Fig. 2D-E) suggested molecular determinants of fragmentation propensities. We thus first analyzed all high-quality *O*-glycan spectra juxtaposing composition-matched glycans containing GalNAcα1-3 or GlcNAcβ1-3, confirming systematic fragmentation propensities on a global scale (Supplementary Fig. 12).

### Molecular dynamics reveals mechanistic basis for diagnostic fragmentation behavior

In the abovementioned scenario (Fig. 2D-E), our conclusion was that GlcNAcβ1-3(Siaα2-6)GalNAc fragmented along the HexNAc-HexNAc axis, while GalNAcα1-3(Siaα2-6)GalNAc fragmented along the Sia-HexNAc linkage. To elucidate how structural properties of these molecules could give rise to these differences in fragmentation behavior, we engaged in molecular dynamics simulations of both isomers.

The fragmentation pattern of the GlcNAcβ1-3(Siaα2-6)GalNAc glycan displayed evidence of a charge induced fragmentation mechanism (Fig. 2D-E). In agreement with this, we saw evidence of the carboxylic acid moiety of the terminal sialic acid interacting with the hydrogen of the C6 hydroxyl group of the terminal HexNAc sugar (Fig. 3). The interaction was sampled 11.9% of our cumulative 2 μs simulations of GlcNAcβ1-3(Neu5Acα2-6)GalNAc. As these simulations were conducted in aqueous solution, rather than a vacuum as would be the environment for fractionation, the frequency of this interaction will be far greater during the *in vacuo* fragmentation due to absence of water molecules competing for hydrogen bonding. Therefore, this suggests that the charge induced fragmentation mechanism of GlcNAcβ1-3(Neu5Acα2-6)GalNAc is due to removal of a proton from the terminal HexNAc sugar, therefore resulting in fragmentation along the HexNAc-HexNAc axis.

**Figure 3.**
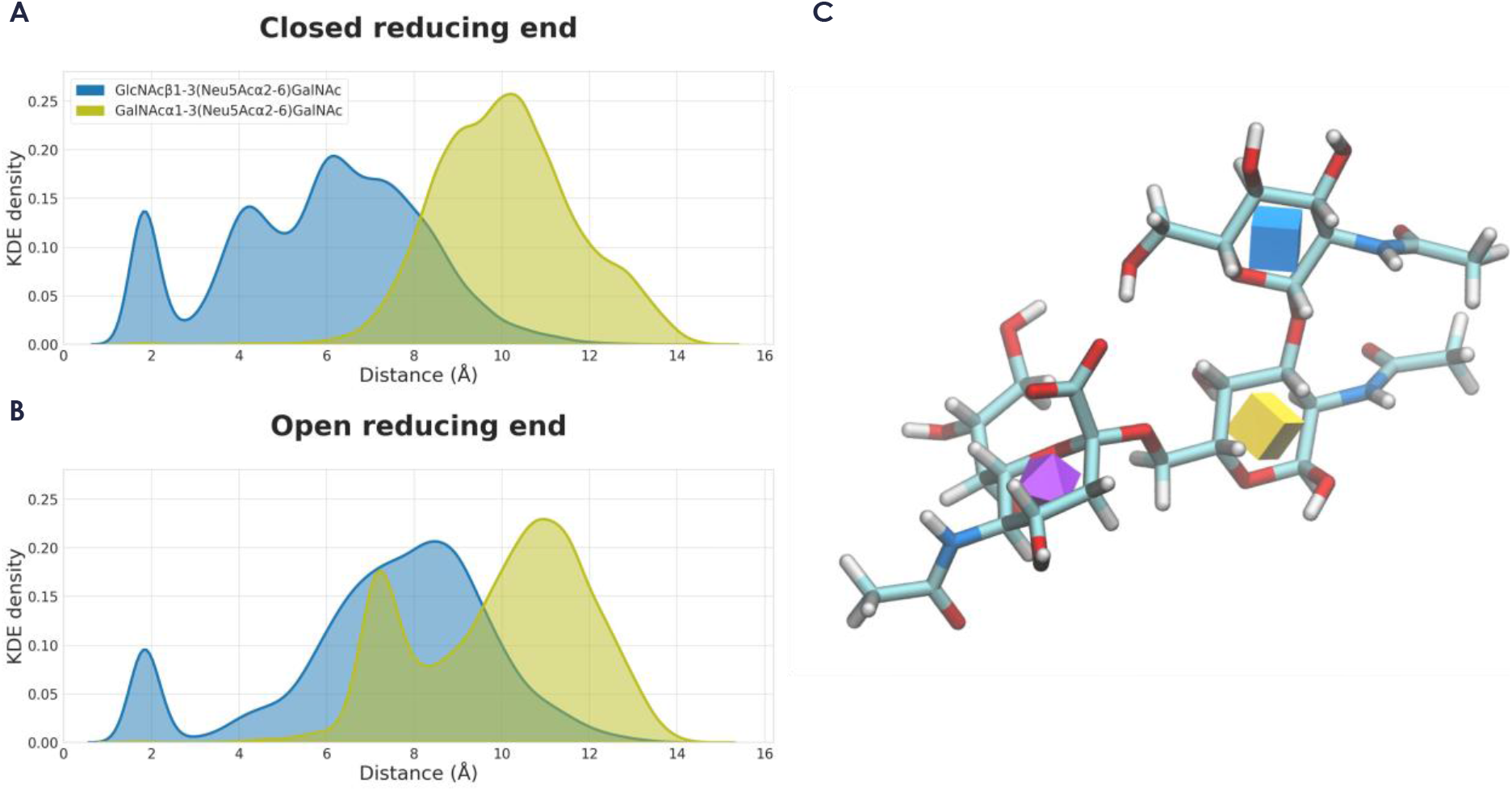
Molecular dynamics reveals fragmentation mechanism. **A-B)** Kernel density estimate distribution of the distance between the center of geometry of the carboxyl group of the sialic acid and the hydrogen of the hydroxyl group of C6 of the terminal HexNAc residues for the closed (A) and open (B) reducing GalNAc residue for both GlcNAcβ1-3(Neu5Acα2-6)GalNAc (blue) and GalNAcα1-3(Neu5Acα2-6)GalNAc (yellow). The plots show how in GlcNAcβ1-3(Neu5Acα2-6)GalNAc, the carboxyl group is able to interact with the hydroxyl of the C6 of the HexNAc. However, this interaction is not observed in GalNAcα1-3(Neu5Acα2-6)GalNAc. **C)** Representative snapshot of the structure of GlcNAcβ1-3(Neu5Acα2-6)GalNAc. A representative snapshot of the structure of GlcNAcβ1-3(Neu5Acα2-6)GalNAc is shown (C), with the interaction between these two moieties displayed a dashed line (orange).

Conversely, simulations of GalNAcα1-3(Neu5Acα2-6)GalNAc were not able to sample this interaction (occurrence < 0.1%). As a result, fragmentation of this glycan occurs along the Neu5Ac-HexNAc linkage instead.

Furthermore, during the ionization of both of the glycans, reductive β-elimination would result in the reducing end GalNAc being reduced to an alditol. As this linearized structure may result in increased flexibility, we also conducted molecular dynamics simulations of both glycans with a linearized reducing GalNAc. These simulations yielded a similar insight to those described previously. In the reduced GlcNAcβ1-3(Neu5Acα2-6)GalNAc glycan, the carboxyl group of the terminal sialic acid interacted with the hydrogen of the C6 hydroxyl group of the terminal HexNAc sugar during 6.8% of the simulated time. Again, the reduced GalNAcα1-3(Neu5Acα2-6)GalNAc was not able to sample this interaction (occurrence < 0.1%).

We therefore concluded that the identified fragmentation behavior can be used to distinguish between these two isomers, an endeavor that is otherwise challenging without specific enzymatic digestion. This implied that we could use our CandyCrunch & CandyCrumbs - powered approach to distinguish very close structural isomers based on diagnostic fragmentation behaviors, beyond single diagnostic ions or ratios and more akin to how human experts would distinguish them.

### Uncovering new biological insights using CandyCrunch and CandyCrumbs

Striving towards AI-assisted glycomics, we propose our platform as a means to enhance human analysts by (i) saving time, (ii) making annotations more robust, and (iii) analyzing samples more comprehensively. We illustrate the latter point with *de novo* predictions of murine intestinal glycans that were too low in abundance to be included in the original annotation but revealed, e.g., the presence of Neu5Gc-containing glycans and low levels of sialyl-Tn antigen in these samples (Supplementary Fig. 13). Importantly, we do not claim that human analysts could not have annotated these structures in principle, but rather that very real time- and resource-constraints make this frequently infeasible in practice. This limitation is lifted by CandyCrunch.

To demonstrate that we could apply our developed methods to truly novel samples, we analyzed the serum *N*-glycome of southern bluefin tuna (*Thunnus maccoyii*), which was measured within GPST000182^36^ but never reported in an annotated manner. This resulted in over 50 glycans, including high-mannose, hybrid, and complex structures, with features such as bisecting GlcNAc, core and antenna fucosylation, Neu5Gc, and multi-antennae *N*-glycans (Fig. 4A, Supplementary Fig. 14). In our comprehensive database within glycowork, not a single glycan from *T. maccoyii* has been reported so far, demonstrating that these pipelines can facilitate new discoveries.

**Figure 4.**
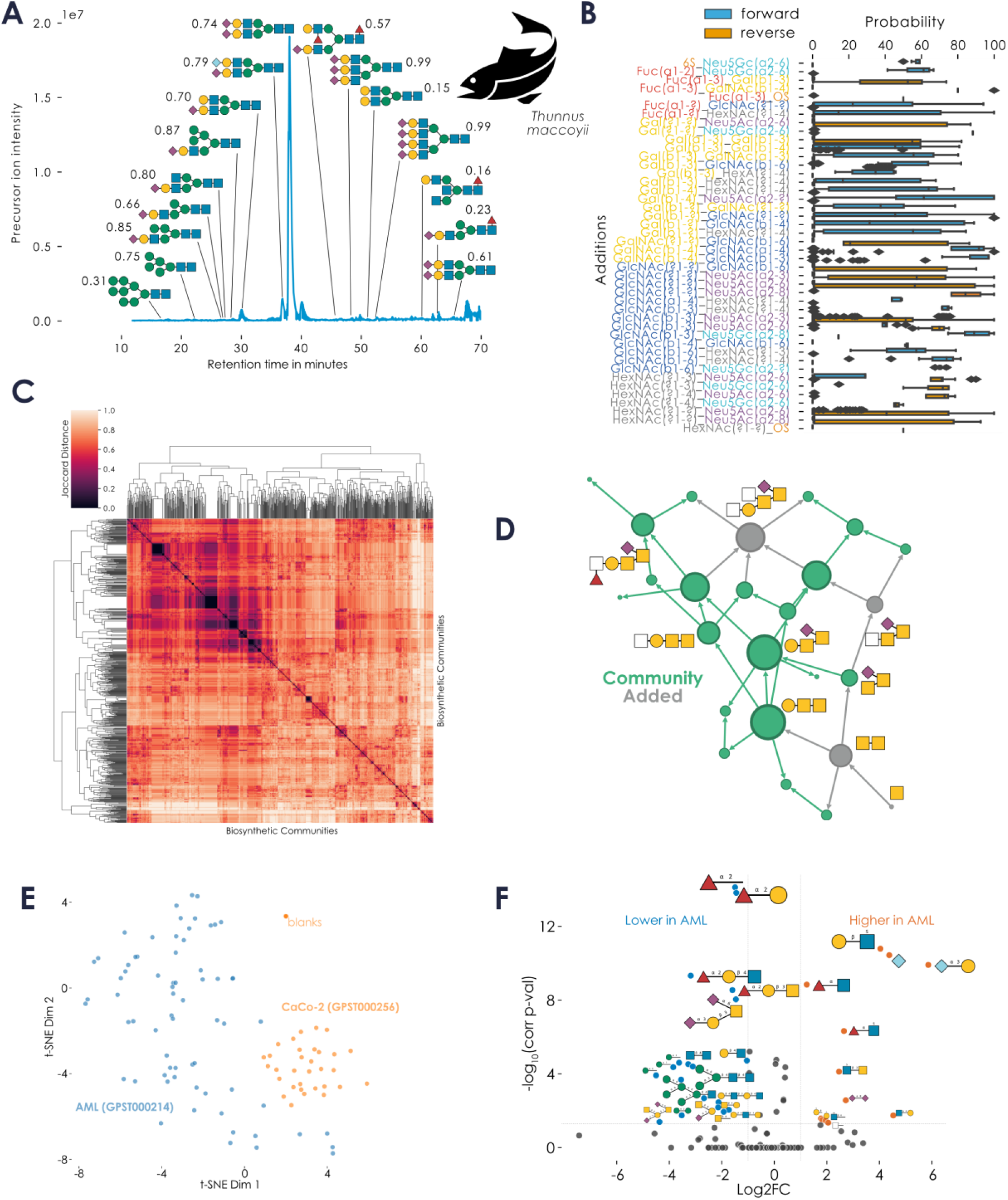
Deriving biological insights from CandyCrunch predictions. **A)** Serum *N*-glycome of the southern bluefin tuna (*Thunnus maccoyii*). Shown are the precursor ion intensities, arrayed by liquid chromatography retention time. Representative structures that are meant to illustrate the identified sequence diversity are shown via the SNFG. Next to each structure, we show the cosine similarity of the shown spectrum and the averaged spectrum of all negative ion mode spectra of reduced glycans of the predicted structure with a confidence above 0.5 (see Fig. 2 for background). **B)** *O*-glycan reactions are path-dependent. For every situation in which two glycosyltransferases competed for the same substrate, we analyzed which order of reactions was experimentally observed across our networks. **C)** *O*-glycan networks decomposed into biosynthetic communities relating to core structures. We detected communities via the Louvain method and calculated their pairwise Jaccard distances, shown here as a hierarchically clustered heatmap. **D)** Community corresponding to core 5 *O*-glycans. Clustering of the distance matrix from (C) using OPTICS (Ordering Points To Identify the Clustering Structure)^38^ resulted in conserved communities broadly corresponding to *O*-glycan cores, with the one from core 5 being shown here as a network, nodes scaled by degree. **E)** Clustering cancer cell line *O*-glycomes. Predicted *O*-glycomes of Acute Myeloid Leukemia (AML) cell lines (GPST000214) and differentiated colorectal cancer cell lines (CaCo-2, GPST000256), via a CandyCrunch model not trained on these datasets, are shown via t-SNE (n = 103), using glycan abundance as features. **F)** Differential glycan expression between AML and colorectal cancer cell lines. Given the predicted glycomes of (E), we used the *make_volcano* function from glycowork to test differential expression at the motif level, shown as a volcano plot. Differentially expressed glycans are drawn inversely scaled by corrected p-value.

We also wanted to highlight how predictions could be used downstream to derive new insights from aggregating glycomics studies. This can even be done in the context of already performed glycomics experiments, distilling results from the accumulated data of many years of study. For this, we re-used the total 250,000 *O*-glycan spectra mentioned in the context of Fig. 2, to construct biosynthetic networks^7^. As described above, this process filled in the gaps of unobserved intermediates in the biosynthesis of observed structures. A key benefit here is that all datasets have been analyzed by the same annotator (CandyCrunch), eliminating an important source of heterogeneity^37^. Applied to our dataset, this resulted in 1,003 biosynthetic networks (corresponding to 1,003 glycomics experiments measuring *O*-glycans) that we used to analyze systematic effects in that glycan class. This revealed that some intermediates were never measured (Supplementary Fig. 15, Supplementary Table 4), such as the reducing end GalNAc (likely due to the mass range of the used mass spectrometer), while others, such as Gal3Sβ1-3GalNAc, were nearly always reliably measured whenever larger structures that included this building block as a substructure were present in a sample. We believe that this approach might shed light on subsets of the *O*-glycome that are currently hard to measure, as we here, again^7^, noted the peculiar absence of GlcNAc-terminated structures from measured glycans as a trend.

Further analyses across our networks then allowed us to compare the reaction order of glycosyltransferases, reinforcing the highly dominant nature of galactosyltransferases^7^ (Fig. 4B). Decomposing the biosynthetic networks into communities unveiled several conserved clusters that were modular and occurred in many of our datasets (Fig. 4C). Further investigation resulted in the observation that these clusters corresponded to the *O*-glycan core structures and their respective biosynthetic extensions (Fig. 4D). In general, these proved to be relatively modular, except for cases such as core 1 and 2, which showed some biosynthetic overlap. We envision that this rapid decomposition of many networks into biosynthetic subcategories will prove useful for comparing and understanding the eventual terminal motifs that will be exposed in these different *O*-glycan cores, as well as their biosynthesis.

As a proof of concept, to demonstrate the capabilities of CandyCrunch for high-throughput analysis, we next predicted the *O*-glycomes of Acute Myeloid Leukemia (AML) cell lines (GPST000214^39^) and differentiated colorectal cancer cell lines (CaCo-2, GPST000256^40^). With a total of 103 glycomics raw files for this analysis, we could show that the predicted glycomes of AML and colorectal cancer cell lines formed distinct clusters (Fig. 4E), which both were separate from the blanks used in GPST000256. We then engaged in a differential glycan expression analysis to investigate what distinguished these clusters. While there was considerable intra-cluster heterogeneity, this analysis revealed that the colorectal cell lines on average were more enriched in structures containing fucosylated galactose and remnant *N*-glycans, while the AML cell lines exhibited higher levels of sialylated glycans and Lewis structures (Fig. 4F). This set of analyses shows that CandyCrunch can be applied to large sets of glycomics measurements and eventually be used in conjunction with other glycowork functionality to reveal dysregulated glycans and glycan motifs, directly from LC-MS/MS raw files.

## Discussion

We present here generalizable methods to (i) predict glycan structures from LC-MS/MS data using deep learning (CandyCrunch) and (ii) automatically annotate fragment ions in MS^n^ spectra (CandyCrumbs). Both CandyCrunch and CandyCrumbs are suited for high-throughput usage and can scale to large datasets as well as extremely diverse glycans and experimental set-ups. With the high performance that we demonstrate here, we are confident that these pipelines will be useful both for experts, accelerating and augmenting their workflows, as well as for less experienced users, similar to how automated workflows in other systems biology disciplines have democratized access to state-of-the-art methods^41,42^.

Our approach is ultimately limited by the representativeness of available data. While CandyCrunch is applicable to all major glycan classes and most experimental setups, we do note that the very best results can be expected for reduced glycans in negative mode, particularly *O*-glycans or free oligosaccharides. This is both a result of high-quality data in those cases and particular efforts in fine-tuning our pipeline for optimal results, as they intersected most with our own research interests and capabilities. In general, compelling results can be expected for samples similar to our training data, strongly enriched in mammalian and fish samples (Supplementary Fig. 16), and we expect to perform worse, on average, on remote samples such as from invertebrates. We envision that, with increasing data, this will improve. We thus urge the community to make their glycomics data (as well as high-quality annotations) available through platforms such as GlycoPOST^43^, as this will improve approaches such as CandyCrunch, and ultimately advance glycobiology and its applications.

We recognize that, as with any model, CandyCrunch predictions are imperfect, exhibiting false negative and false positive predictions, which occasionally might not resemble errors made by humans. Particularly non-CandyCrunch glycan additions within our pipeline, via biosynthetic networks and database queries, exhibit a more tentative character and should be further evaluated by experts. For ideal results, we always recommend predictions to be further refined by experts. We are, however, convinced that CandyCrunch predictions can raise result quality and comprehensiveness for both experts and novices, in addition to the considerable increase in throughput. Lastly, during data curation, we assumed expert annotations within our training data to be correct, which may retain analyst bias, such as preferential annotation of type II versus type I LacNAc structures in *N*-glycans without conclusive evidence. Once sufficient data become available, future work may extend this approach to higher-order MS^n^ spectra and/or exoglycosidase treatments, with more detailed structural information.

Beyond the fact that the zero-shot capabilities of CandyCrunch are limited, we would also like to note that, while we support common derivatizations such as permethylation, we do not currently support every type of glycan modification within CandyCrunch and CandyCrumbs. Specialized methods, such as azidosugars^44^, are at the moment beyond our scope.

We are enthusiastic about the potential of upcoming methods to simulate high-resolution fragmentation spectra via deep learning^45^, which could be adapted for AI-glycomics in future work and aid either training or the evaluation of prediction results. While we focus on glycomics here, we envision that analogous efforts in glycoproteomics could also advance and accelerate the field. Overall, we conclude that our presented methods not only pave the way for AI-enhanced structural glycomics but also enable many other avenues ranging from systematic comparisons over data science to glycoinformatics. This is facilitated by our large, curated dataset and the ability to quantify spectrum quality, engaging in analyses at scale for many different aspects of glycomics data.

## Online Methods Dataset

Tandem mass spectra stemmed from repositories such as GlycoPOST^43^, MassIVE, UniCarb-DB^33^, UniCarb-DR, and NIST, as well as from individual publications with associated public raw data. A full list of the 189 data sources can be found in Supplementary Table 5. All raw files were converted into the open-access format .mzML using the msconvert software^46^. A custom script using the pymzML package^47^ (version 2.5.2) was used to extract all spectra at the MS/MS level, together with their stored precursor ion m/z and retention time, if available. This extraction functionality is now available as the *process_mzML_stack* function within our CandyCrunch package (version 0.1), next to an analogous *process_mzXML_stack* function. We extracted up to 1,000 fragment peaks of the highest intensity per spectrum, if available. Then, spectra were retained that fell within +/-0.5 Da m/z and +/-2 minutes retention time of reported glycan peaks in the associated publications. All retained spectra were kept for self-supervised training, paired with the information of the respective glycan class, while only spectra that could be unambiguously linked to structures described in the respective publications were kept for supervised training. This resulted in a total number of 625,547 glycan spectra, of which 485,406 spectra were labeled with a defined structure and could be used for training, the latter stemming from 3,508 unique glycan structures (Supplementary Table 6). The full dataset can be found at Zenodo under the doi:10.5281/zenodo.7940047.

### Data Processing

We first removed all spectra with a retention time below two minutes as noise. Retention times then were normalized for each individual sample, by dividing absolute retention times by the respective maximal retention time (or a minimum of 30, if the maximum extracted retention time was below 30). Missing retention times were assigned a value of zero. Fragment intensities were normalized for each spectrum, by dividing the intensity of each peak by the total intensity of the spectrum. Then, intensities were binned in 2,048 equal-sized m/z windows from the observed minimum (39.714) up to a maximum of 3,000. Additionally, the m/z remainder (i.e., the difference of the m/z of the highest intensity peak of a bin to the left bin window) was calculated for each bin, as suggested in Altenburg et al^25^, allowing the model to learn exact peak location despite binning. Glycan class, mass spectrometry ion mode, ion trap type, LC type, and glycan modification type were coded as integers to allow for learned embeddings.

During training, we capped all glycan structures to at most 1,000 randomly sampled spectra per structure, to avoid imbalance by frequently observed but simple glycans. In the self-supervised setting, we preferentially sampled spectra from labeled examples and only supplemented, where needed, by unannotated spectra labeled by the model. This led to a final dataset of 300,363 spectra. We used an 85/15 split into train/test set, which was split on the level of samples, to ensure that spectra of one sample were not found in both train and test and thus make the generalizability estimation more robust. For training, classes in the test set that would constitute zero-shot prediction were afterwards moved into the train set.

### Model architecture

CandyCrunch is a dilated residual neural network, with additional embedded inputs, to predict glycan structure from tandem mass spectrum in a multiclass classification setup.

For the processing of binned intensities and m/z remainders, a 1D-convolution layer was followed by a leaky ReLU and six residual dilated convolutions, with dilations of 1, 2, 4, 8, 16, and 32. Then, we used max-pooling with a kernel size of 20 and a fully connected layer to bring this output to a dimensionality of 1,024. Glycan class, mass spectrometry ion mode, ion trap type, LC type, and glycan modification type were embedded into dimensionalities of 24, respectively. Precursor m/z and normalized retention time were also brought to dimensionalities of 24 via a fully connected layer, a layer normalization, and a leaky ReLU. Then, all inputs were concatenated and passed through two sets of fully connected layers, layer normalization, leaky ReLUs, and dropout (at a rate of 0.2). Finally, a last fully connected layer yielded the class probabilities. In total, CandyCrunch exhibited 12,375,084 trainable parameters.

### Model training

All models were trained in PyTorch^48^ (version 1.13.1) using two Zotac GeForce RTX 4090 Trinity GPUs. CandyCrunch was initialized via He initialization. All models were trained for 200 epochs, with an early stopping regularization of stopping training after 12 epochs without improvement in the test loss and a batch size of 256.

We set the learning rate at 0.0001, with a schedule to reduce the learning rate to a fifth after four epochs with no improvement in test loss. As a base optimizer we used AdamW with a weight decay of 2e-5, which was further modified via Adaptive Sharpness-Aware Minimization (ASAM)^49^ to ensure a generalizable final model.

Data augmentation during training was used only on the training set and included random (i) low-intensity peak removal, (ii) peak intensity jitter, and (iii) new peak addition for the binned spectrum, as proposed previously for mass spectrometry^50^, as well as adduct formation of the precursor ion (acetate/sodium adducts) and random noise of the precursor m/z (+/-0.5 Da) and retention time (+/-10%).

As our base loss, we used PolyLoss^51^, with an additional label-smoothing of 0.1 and epsilon = 1. We also used two additional loss terms, informed by domain knowledge, that were added to the PolyLoss term. These constituted a structure distance loss and a composition distance loss. Both involved the calculation of a distance matrix, based on pairwise cosine distances of fingerprint vectors of either the number of mono- and disaccharide motifs or the base composition of two glycans. All operations on glycans were performed using glycowork^31^ (version 0.7). Then, the class probabilities for each input sample, transformed via a softmax activation, were multiplied by the structure distance vector and the composition distance vector (i.e., the distance to the target glycan), followed by mean averaging to obtain loss terms. This unsupervised procedure preferentially penalizes confidently predicted but structurally dissimilar glycans and improves performance as well as the meaningfulness of errors.

We first engaged in supervised training on annotated MS/MS spectra. Then, using the trained model we predicted glycan structure for our unannotated spectra for self-supervised training. Spectra with a prediction score of over 0.7 were then merged with the original training dataset, followed by a deduplication step. Specifically, as described above, we retained the same test set and again formed a training dataset with at most 1,000 examples per glycan, with preferential sampling from the original dataset and only supplemental sampling from the extended dataset, followed by re-training.

### Model inference

To predict glycan structures from unannotated raw files, all tandem spectra were extracted via pymzML as described above and processed as described for the general data processing. Then, we grouped m/z precursor ions by scanning for discontinuities larger than 0.5 Da in the extracted spectra. Within these m/z groups, we searched for structural isomers by analyzing their retention time, scanning for discontinuities above 0.5 minutes. For each retention time group, we averaged all spectra for input of a robust averaged spectrum to CandyCrunch and extracted the median spectrum, to have a representative spectrum for each glycan entity in the sample. We first retrieved the top 25 predictions for each averaged spectrum, using the trained CandyCrunch model. We then employed a single-parameter variant of Platt Scaling^34^ to calibrate the prediction confidence prior to the softmax layer, using a scaling factor of 1.15 that was estimated via the L-BFGS algorithm. Using test-time augmentation, we averaged the predictions of five independent inferences that were modified with the same data augmentation strategy as employed during training.

Next, we used domain knowledge to automatically filter out predictions, such as of (i) a prediction probability below a threshold of 0.01, (ii) the wrong glycan class, (iii) the wrong mass, even when considering multiply-charged ion forms, and (iv) predictions that lacked corroborating diagnostic ions in their fragment lists. Domain-specific exceptions were made, such as allowing cross-class predictions if the prediction confidence was extraordinarily high (above 0.2; justified by the fact that *O*-linked glycan samples often contain remnant *N*-linked glycans etc.) Finally, predictions were deduplicated by merging any mass / retention windows that resulted in identical predictions.

Lastly, we used biosynthetic knowledge to refine our predictions, conceptualized in the *canonicalize_biosynthesis* function within CandyCrunch. Using the *subgraph_isomorphism* function from glycowork and starting from the largest glycan prediction, we searched for top1 predictions of biosynthetic precursors in the whole prediction dataframe. For each prediction at mass M, we added 0.1 to its prediction confidence for each unique biosynthetic precursor in top1 predictions at mass M-1, M-2, … M-n. If this changed the order of predictions, we reordered predictions according to their score. Thereafter, scores were re-normalized to 1 and the, up to, top5 predictions were retained. This procedure not only improved the accuracy of our results but also increased the meaningfulness and consistency of both correct and wrong predictions (i.e., wrong predictions were structurally closer to the ground truth after this procedure).

Spectra without valid predictions but with valid compositions, cross-referenced by relevant databases within glycowork, were also retained and subjected to as many of the abovementioned domain filters as possible. The whole inference workflow, including elements described below, is available via the *wrap_inference* function in the CandyCrunch package.

### Zero-shot prediction

For a given sample, all retained top1 predictions were used to construct a biosynthetic network as described previously^7^, using the implementation within glycowork. For milk oligosaccharides, this also included evolutionary pruning, as pre-calculated species networks were available. Then, we calculated whether any of the inferred biosynthetic precursors would explain the mass and composition of glycan spectra without a valid prediction. Matches within a mass difference of 0.5 Da, including multiply-charged ions, were retained as additional predictions beyond our model-defined library of predictable glycans. While direct model predictions were awarded the evidence category “strong”, the biosynthetic network intermediaries merited the category “medium”.

Next, we checked for missed Neu5Gc-substituted Neu5Ac-glycans and vice versa (i.e., a mass difference of 16 Da, with the corresponding diagnostic ions). Additionally, we used a suitable subset of the glycowork-stored database, of the right taxonomic section and glycan class, to search for possible matches to compositions without predictions. Both of these endeavors were annotated with the evidence label “weak”.

After these additional routines to enable predictions outside of our defined list of glycans, we again employed the domain-knowledge informed filters mentioned above. This ensured that glycans introduced via these methods still had empirical support in the underlying data. Predictions from these routines were also subjected to the *canonicalize_biosynthesis* workflow from above (though “bonus” points were only awarded for biosynthetic precursors from the “strong” category), to allow for prioritization of the most probable structures.

### Fragment annotation via CandyCrumbs

The final prediction of the CandyCrunch model was used as a starting point for fragment annotation and converted into a directed graph using NetworkX (version 3.0), each monosaccharide making up a node and each linkage labeling an edge. The randomized enumeration method was implemented to find all induced connected subgraphs^52^. After filtering which modifications are physically possible based on linkage numbers, each terminal monosaccharide on the subgraphs was permuted to create these cross-ring or bond fragmentations. Each possible global modification was also added to each fragment. The mass of each theoretical fragment was calculated to then be matched with observed masses in MS^2^ spectra. Finally, the fragments were converted into Domon-Costello^32^ and IUPAC-condensed nomenclature. If multiple fragment possibilities could explain a given m/z value, a prioritization scheme was developed (Supplementary Fig. 9), which emphasized prior likelihood of each fragment option and the evidence of the remaining fragments in a given tandem spectrum. CandyCrumbs is available via *CandyCrunch*.*analysis*.*CandyCrumbs* in our developed Python package.

### Molecular dynamics simulation

Initial conformations for the GlcNAcβ1-3(Neu5Acα2-6)GalNAc and GalNAcα1-3(Neu5Acα2-6)GalNAc glycans were obtained using the Carbohydrate Builder tool of the GLYCAM-Web server^53^. Four structures were produced for each glycan with different combinations of the α2-6 torsion angles. This approach provided different initial starting points for the simulations, and thus maximized the sampling of the conformational space. Each glycan was parameterized with the GLYCAM06-j1 force field^54^, and a cuboid solvent box of TIP3P water molecules created to produce a minimum solute distance of 15 Å. In the case of the reduced glycan structures, the structures of the open GalNAc were parameterized using the GAFF2 forcefield^55^. A single Na^+^ ion was included in each system to neutralize the net charge of the system. These systems were then converted into GROMACS topology files using Acpype^56^. For each initial starting conformation of each system, a 500 ns simulation was performed using GROMACS2022.4^57^, resulting in 2 μs of simulations for each respective system.

### Biosynthetic network analysis

For all networks constructed and analyzed in this work, we used the code functionality within the glycowork.network.biosynthesis module (version 0.7). Our analyses were oriented very closely by the ones described in Thomès et al.^7^ Briefly, the analysis of glycosyltransferase competition was performed by analyzing diamond-like network motifs via the *trace_diamonds* and *find_diamonds* functions within glycowork. Thereby, we analyzed the proportion of networks that presented a certain case of glycosyltransferase competition and counted how often each alternative order of reactions was experimentally observed among these. This allowed us to analyze which reaction order dominated across (i) glycan contexts and (ii) networks. The differences shown in Figure 4 were further filtered to contain at least (i) two glycan sequence contexts, (ii) a mean difference of 30, and (iii) a corrected p-value below 0.01.

Biosynthetic communities were extracted using the *get_communities* function, from glycowork, on reaction path preference-pruned biosynthetic networks^7^. Conserved communities were detected by first calculating a distance matrix based on pairwise Jaccard distances, followed by clustering these distances using the OPTICS algorithm as implemented in scikit-learn (version 1.2.2), with a minimum number of 50 samples per cluster.

### Statistical analyses

Comparing two groups was done via one-tailed or two-tailed Welch’s t-tests. In all cases, significance was defined as p < 0.05. All multiple testing was corrected with a Holm-Šídák correction. All statistical testing has been done in Python 3.9 using the statsmodels package (version 0.13.5) and the scipy package (version 1.10.1). Effect sizes were calculated as Cohen’s d using glycowork (version 0.7). The correlation of distance matrices was performed via two-sided Mantel tests as implemented within scikit-bio (version 0.5.8).

## Data and Code availability

All relevant code is integrated into glycowork (version 0.7) and/or can be found at https://github.com/BojarLab/CandyCrunch. CandyCrunch and CandyCrumbs can also be readily accessed at https://colab.research.google.com/github/BojarLab/CandyCrunch/blob/main/CandyCrunch.ip ynb. All relevant data can be found at Zenodo under the doi:10.5281/zenodo.7940047 or is contained within the supplementary material.

## Acknowledgments

This work was funded by a Branco Weiss Fellowship – Society in Science awarded to D.B., by the Knut and Alice Wallenberg Foundation, and the University of Gothenburg, Sweden. The Science Foundation of Ireland (SFI) Frontiers for the Future Programme is gratefully acknowledged for financial support of CMI (20/FFP-P/8809). We also thank Carl Fogarty for his assistance in parameterizing the open GalNAc residue for molecular dynamics simulations.

## Author Contributions

D.B. and J.U. conceived the method. D.B. curated the dataset. D.B., E.F., C.M.I., and J.U. performed computational analyses. D.B., C.M.I., and J.U. prepared the figures. C.J., N.G.K., and K.A.T. confirmed spectra annotations and provided domain-expertise for method development and application. D.B. and E.F. supervised. All authors wrote and edited the manuscript.

