## Supplemental Figures for "Predicting glycan structure from tandem mass spectrometry via deep learning"

**Supplementary Figures**

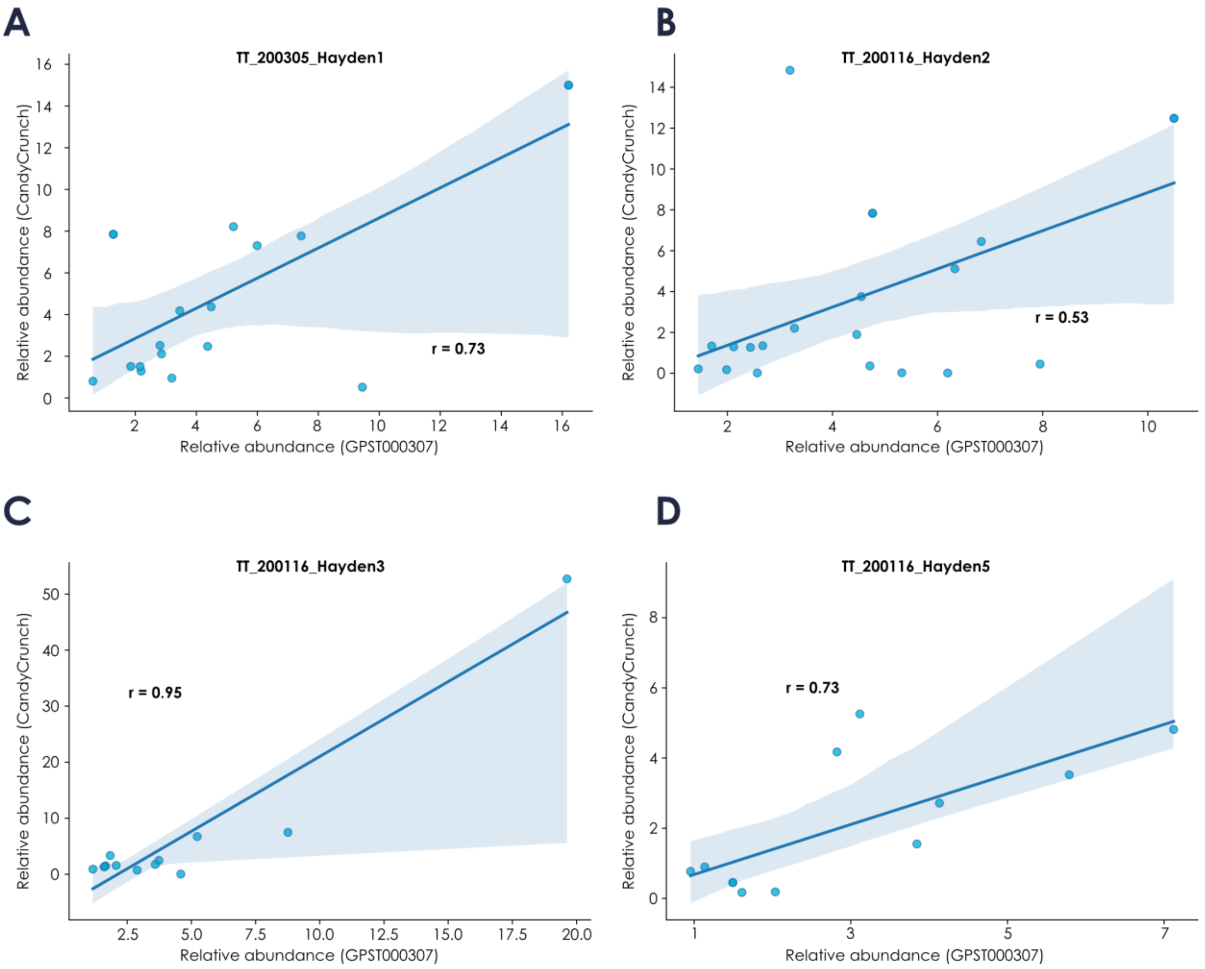


**Supplementary Figure 1. Relative abundances obtained by CandyCrunch correlate with relative abundances from experts. A-D)** For the four *O*-glycan raw files from GPST000307 (GlycoPOST), we extracted the ion intensity from each precursor selected for fragmentation. Then, we used CandyCrunch to predict glycan structures and paired those with relative abundances (in percent) estimated from the ion intensities. In each sample, for those glycans for which we could unequivocally establish equality between expert annotation and prediction, we compared their relative abundances quantified by the analyst as well as CandyCrunch. All correlations between experimental data and predictions were done via fitting a linear regression and r represents Pearson's correlation coefficient.

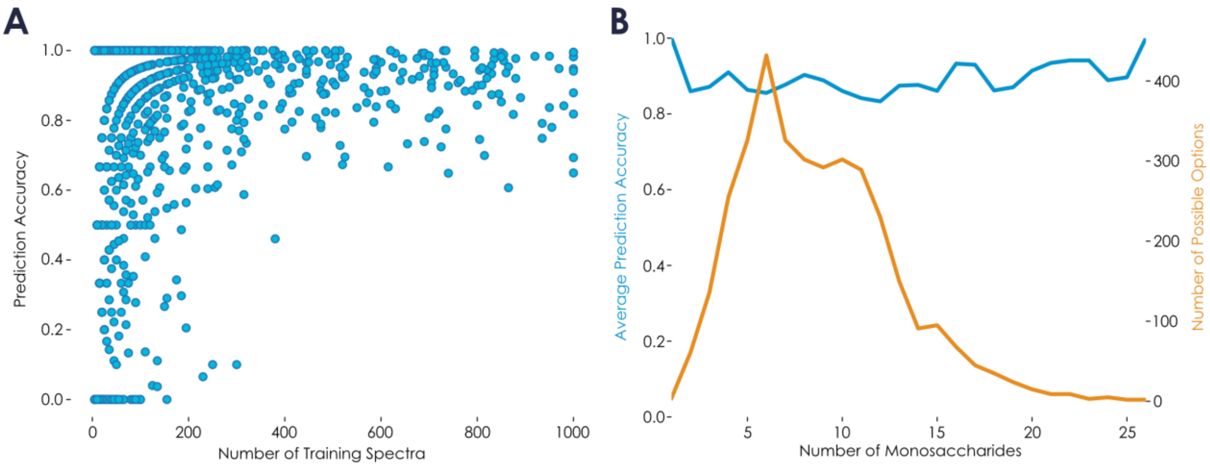


**Supplementary Figure 2. Dependence of predictive performance on number of example spectra and size for each glycan. A)** For all 1497 unique glycans occurring in the independent test set, the prediction performance of CandyCrunch on that glycan in the independent test set is plotted against the number of example spectra of that glycan in the training set. The plot shows that CandyCrunch exhibits high predictive performance for most glycans and that relatively few example spectra for a new glycan are needed to achieve high predictive performance. **B)** Similarly, we plotted the average prediction per glycan size (i.e., number of monosaccharides) in the independent test set. While this shows a stable prediction accuracy across glycan sizes, we note that the number of possible options (i.e., how many glycans of size 26 are in our dataset) drastically decreases with glycan size, suggesting that the indicated accuracy for very large glycans might be overly optimistic.

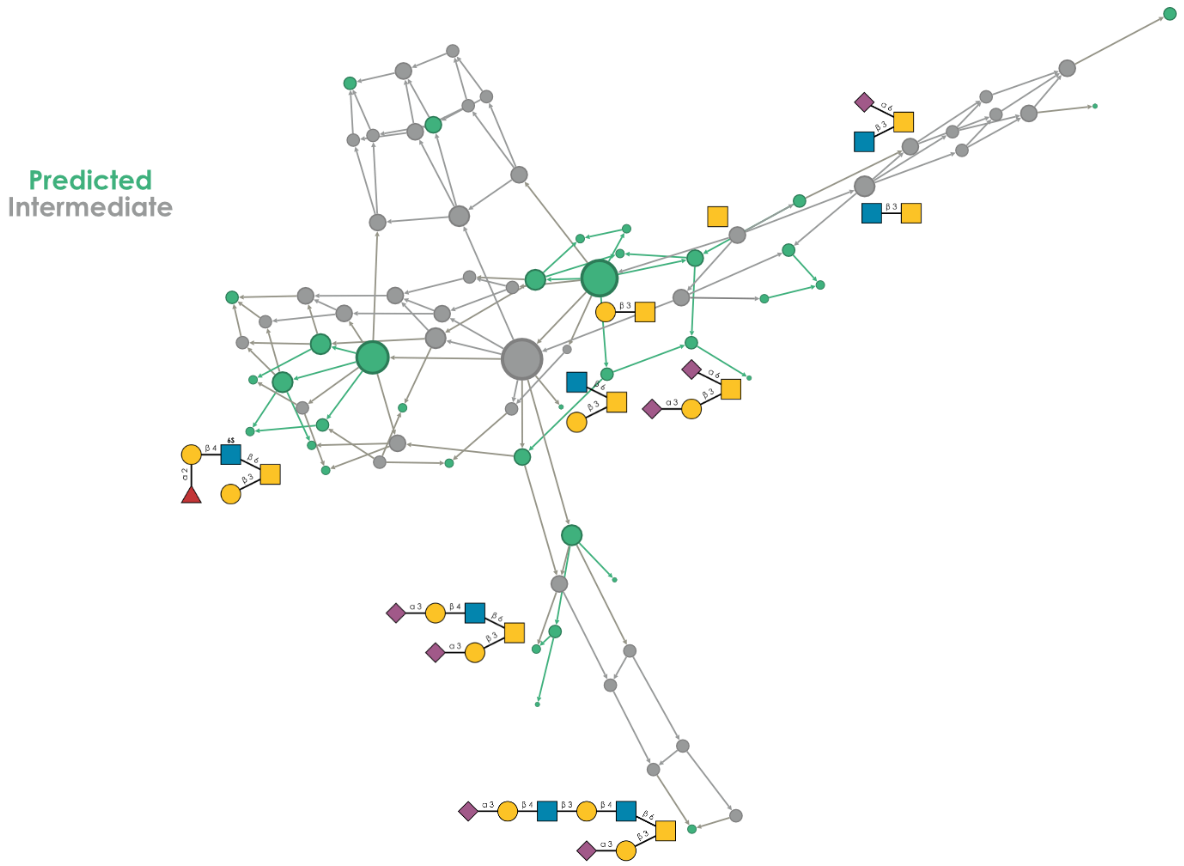


**Supplementary Figure 3. Using biosynthetic networks for interpolative zero-shot predictions.** The biosynthetic network was constructed as described by Thomès et al., 2023, *bioRxiv*, from glycan predictions of the example file KM-O-gly-CaCo-2-900sps-3l-3psi-280-meOH-5D_3_1-0006-0006.mzML from GPST000256. CandyCrunch predictions are colored in green, and inferred intermediates connecting these predictions are colored in gray. Nodes are scaled by degree. Selected structures are drawn via their SNFG representation.

**A**

| | Glycoforest (2017) | GlycoDeNovo2 (2022) | CandyCrunch (ours) |
|---|---|---|---|
| **Approach** | Consensus spectrum network search | Composition constrained topology search | Dilated residual neural network classification |
| **Language** | Java | MATLAB | Python |
| **Runtime** | Minutes | n/a | Seconds |
| **Input** | mzXML | List of peaks | mzML/mzXML |
| **Web hosted** | Yes | No | Yes |
| **Glycan types** | *O*-glycans | *N*-, *O*-, lipid, free glycans | *N*-, *O*-, lipid, free glycans |
| **Stereochemical resolution** | No | Yes | Yes |
| **Linkage type resolution** | No | No | Yes |
| **Supported post-biosynthetic modifications** | Sulfation | - | Sulfation, Phosphorylation, Acetylation, etc. |
| **Experimental setup generalizability** | - | - | Ion mode, liquid chromatography, glycan modification, etc. |

**B**

| | GlycoWorkbench (2008) | glypy (2019) | CandyCrumbs (ours) |
|---|---|---|---|
| **Language** | Java | Python | Python |
| **SNFG Output** | Yes | No | Yes |
| **IUPAC Output** | No | No | Yes |
| **Cross-ring fragments** | Yes | No | Yes |
| **Cross-ring validity checks** | No | No | Yes |
| **Linkage type resolution** | No | No | Yes |
| **Optional fragmentation types** | Yes | No | No |
| **Programmatic integration** | No | Yes | Yes |
| **Fragment prioritization** | No | No | Yes |

**Supplementary Figure 4. Feature comparison between models predicting glycan structure and annotating fragments. A-B)** We compared current state-of-the-art methods for the tasks of predicting glycan structure from MS/MS data (A) and annotating therein resulting fragment ions (B) to assess the capabilities of our CandyCrunch and CandyCrumbs methods, respectively. Representative features were chosen to compare the current strengths and weaknesses of each method as best as possible. Green coloring indicates feature values that were judged to be more desirable. We conclude that, based on these metrics, CandyCrunch and CandyCrumbs present the best-in-class solution to the respective problem.

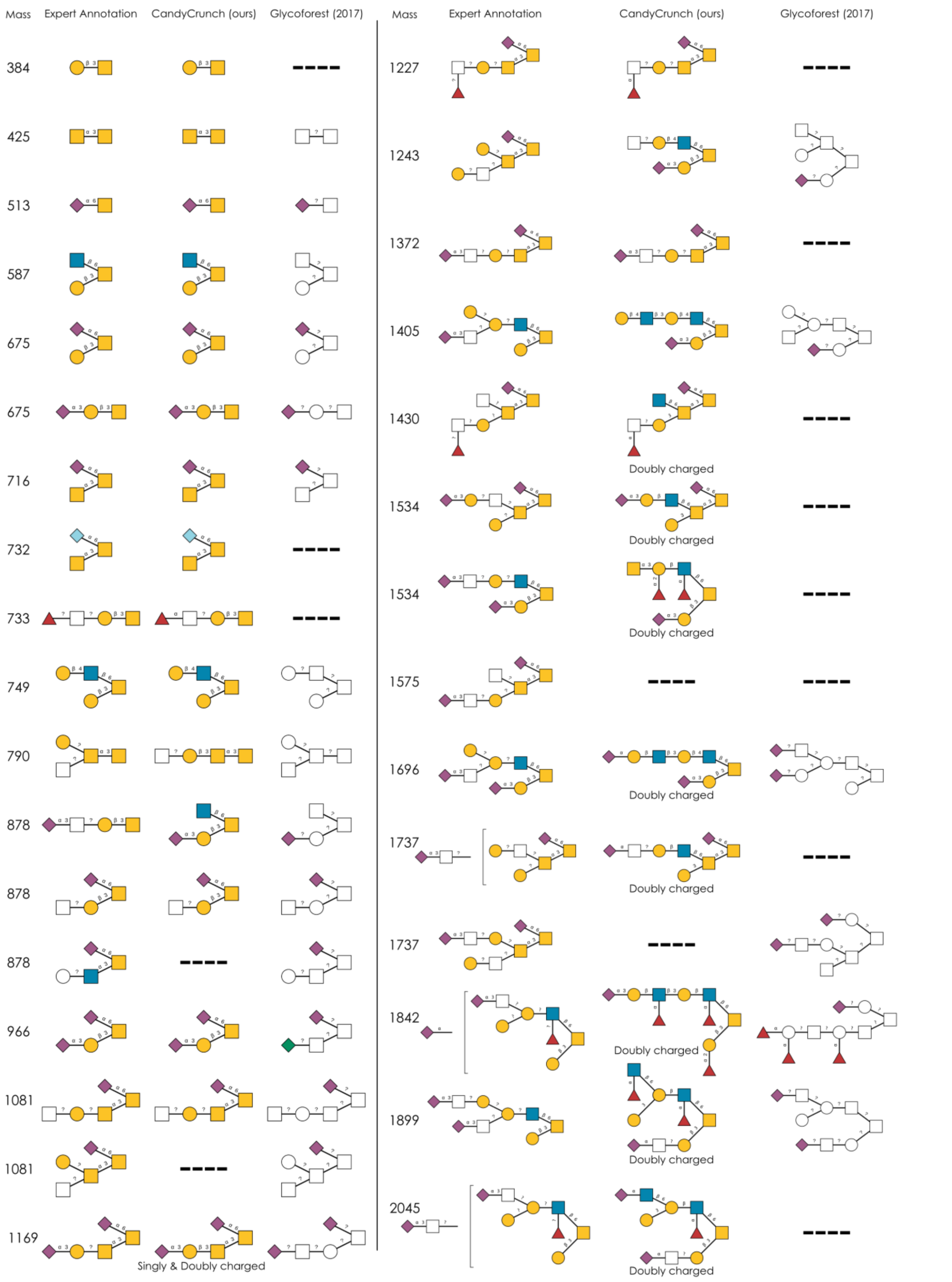


**Supplementary Figure 5. Comparing CandyCrunch with Glycoforest.** For JC_131210PMpx5.mzML, not used for training CandyCrunch but used for developing Glycoforest, we compared predictions by both methods for all expert-annotated structures. Additional CandyCrunch predictions exist beyond expert annotations. Deviations from expert annotations (e.g., 878-1) are not necessarily errors. Glycans are shown via the SNFG, with isomers ordered by retention time.

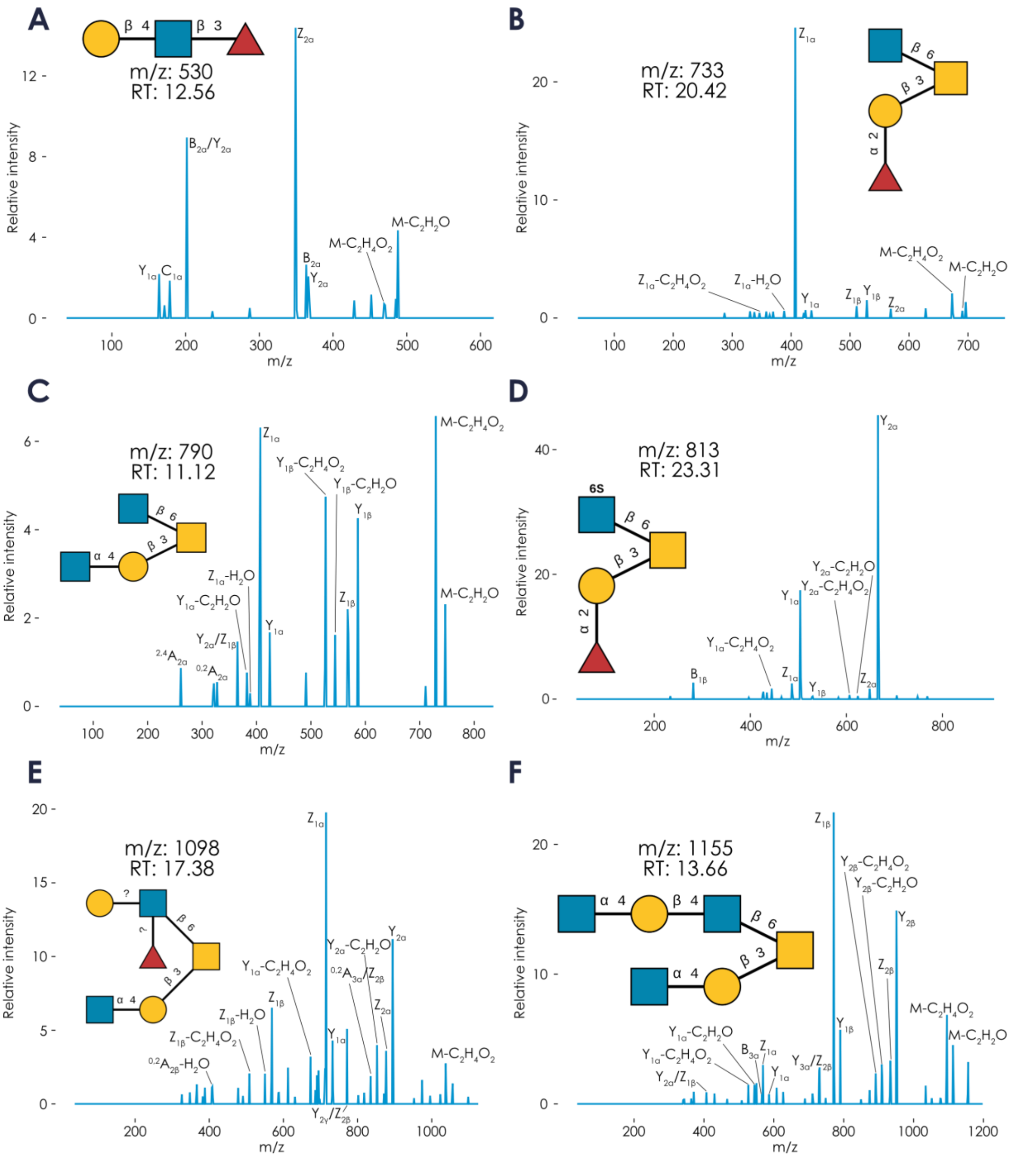


**Supplementary Figure 6. Extending expert annotation with CandyCrunch. A-F)** We predicted all *O*-linked glycan structures from the raw MS/MS file JC_171002Y1 (from Kouka et al., 2022, *Molecules*). Starting from the smallest structures, we chose the first six predicted structures that were not contained in the expert annotation (either new masses or additional isomers at different retention times). We assume that most of these structures, except for (A), based on their abundance and biosynthetic character, could stem from the remnants of the porcine gastric mucin sample used to calibrate the MS instrument, showcasing the exceptional sensitivity of CandyCrunch. The associated MS/MS spectra used for prediction, together with their m/z values, retention times, and predicted structures are shown with their annotated fragments in the Domon-Costello nomenclature.

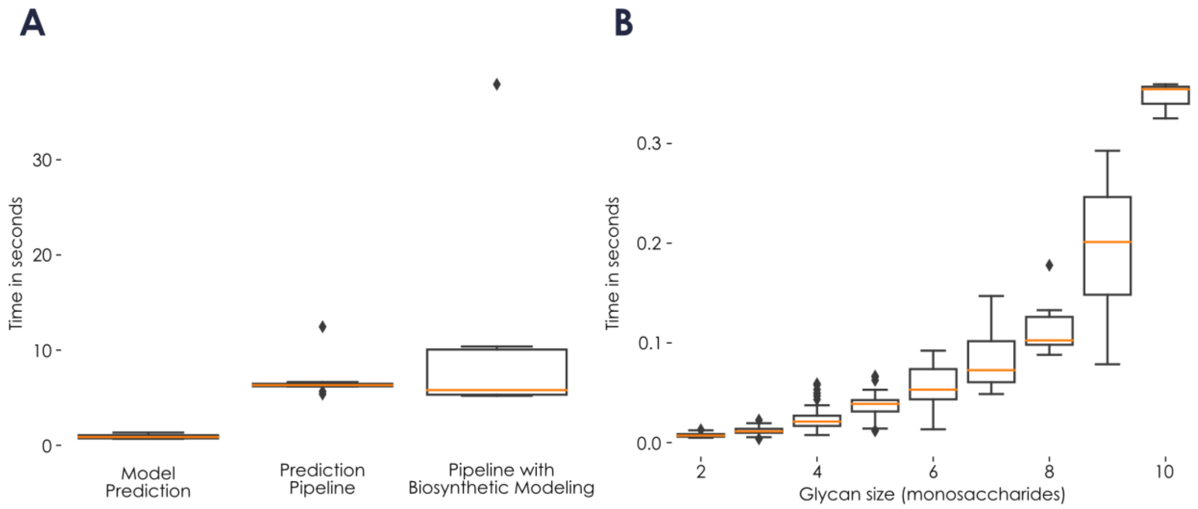


**Supplementary Figure 7. Estimating speed of CandyCrunch and CandyCrumbs. A)** Using all raw MS/MS files from Kouka et al., 2022, *Molecules* (n = 10), we extracted all spectra with process_mzML_stack() and recorded the time in seconds for (i) predicting all spectra with CandyCrunch, (ii) predicting and curating all spectra, and (iii) predicting and curating, including noting spectra with valid compositions but no valid prediction. Predictions were performed using a single Nvidia A100 GPU and the results are depicted via boxplots. **B)** On the same files as in (A), we used all 205 top1 glycan predictions and their spectra to annotate their fragments via CandyCrumbs. Computation time per glycan is plotted against glycan size, using an Intel® Xeon® CPU @ 2.00GHz. Variance within one size can be explained by different degrees of branching.

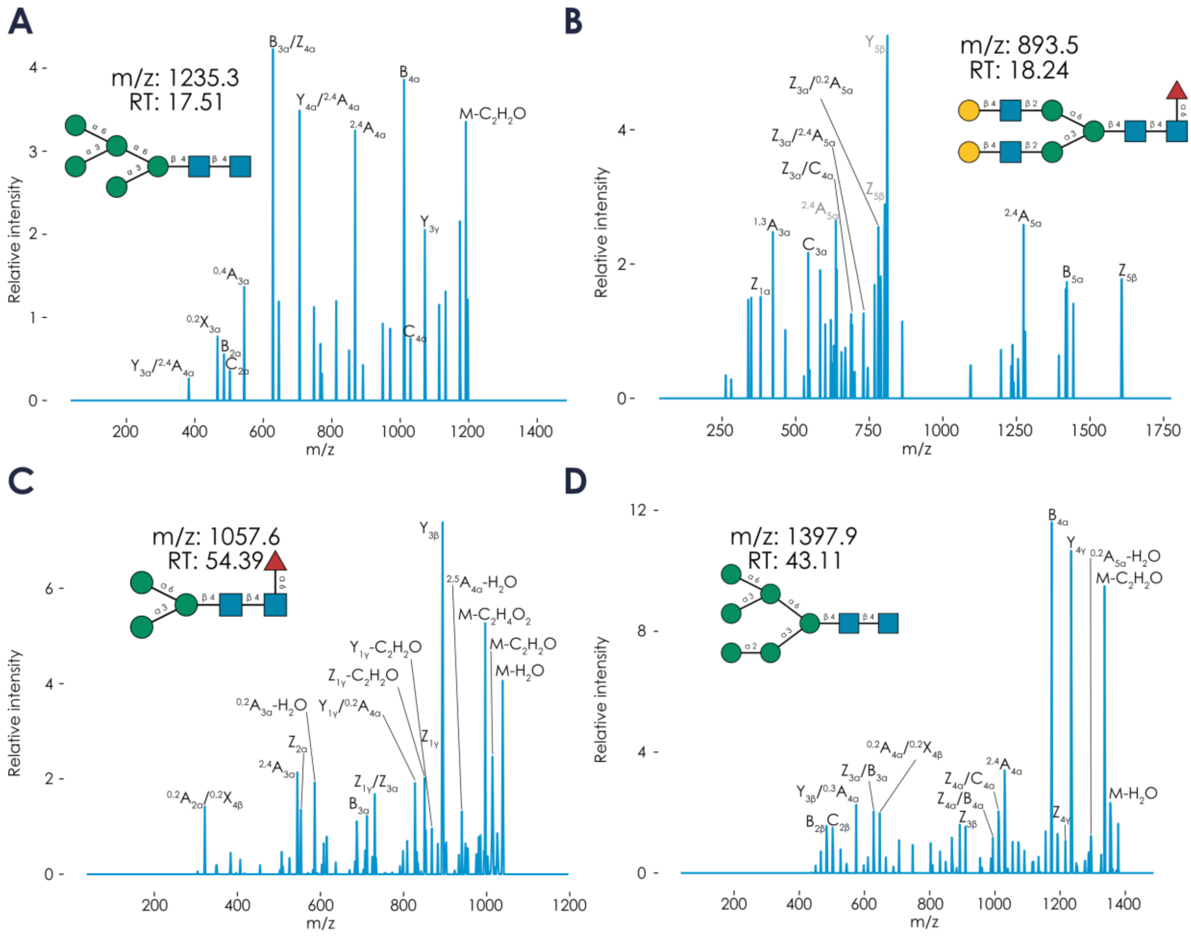


**Supplementary Figure 8. Predicting unexpected glycans. A-D)** Shown are example CandyCrunch predictions of co-released *N*-linked glycans in preparations of *O*-linked glycans that were not annotated in the respective publications. The examples are from JC_171002Y2.mzML (A-B; Kouka et al., 2022, *Molecules*) and KM-O-gly-CaCo-2-900sps-3l-3psi-280-meOH-5D_3_1-0006-0006.mzML (C-D; GPST000256). The associated MS/MS spectra used for prediction, together with their m/z values, retention times, and predicted structures are shown with their fragments, annotated by CandyCrumbs, in the Domon-Costello nomenclature. Doubly-charged fragment ions are denoted with gray text.

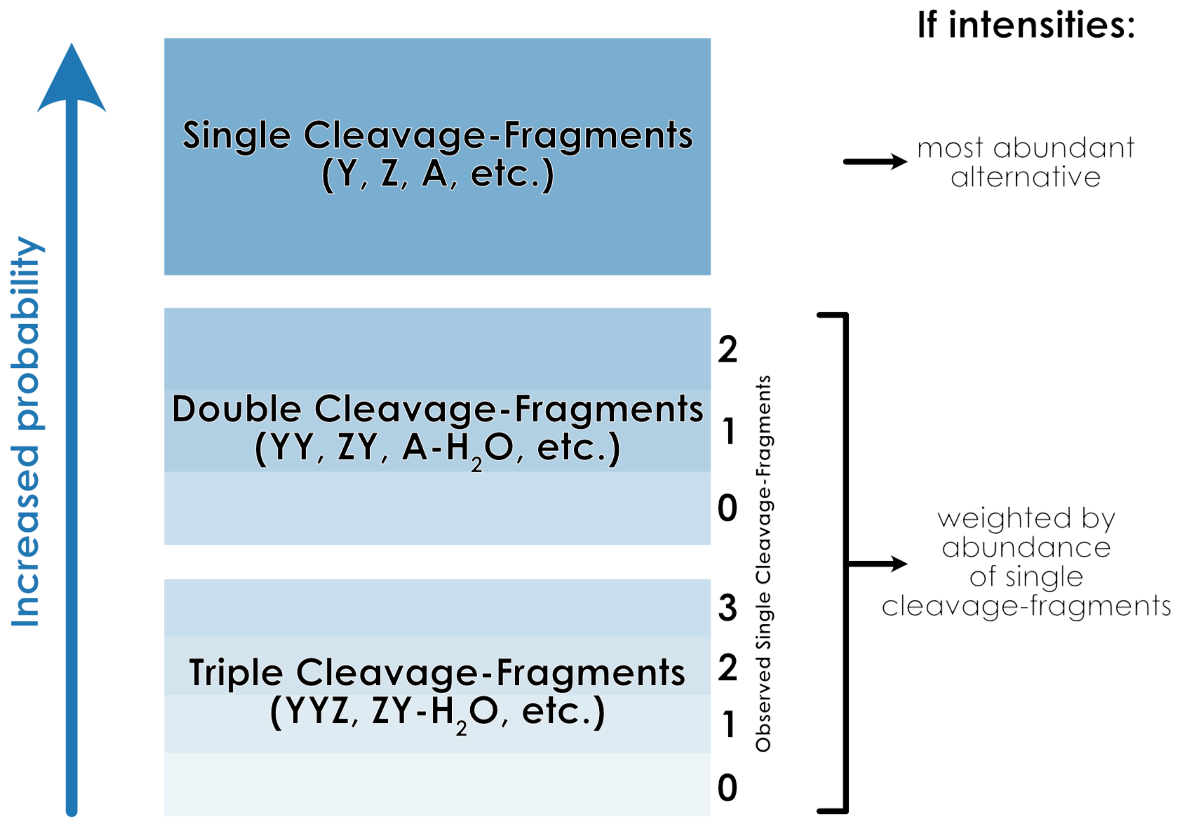


**Supplementary Figure 9. Decision scheme for choosing the best fragment alternative within CandyCrumbs.** From the fragment possibilities, up to triple cleavage-fragments, that explained a given m/z within a specified variance, we selected the most probable explanation via heuristics and domain knowledge. Less complex fragmentation (e.g., single cleavage-fragments) were always preferred over more elaborate fragmentation patterns. Further, evidence of constituent single cleavage-fragment ions (e.g., Y) in a spectrum was interpreted as evidence for the double cleavage-fragment possibilities comprised of these single cleavage-fragment ions (e.g., YZ). If m/z intensities were also supplied, this type of evidence was weighted by the observed abundance of the respective single cleavage-fragments (e.g., Y*intensity(Y)), with the assumption that more prominent single cleavage-fragment ions were also more likely to spawn compound fragmentation products.

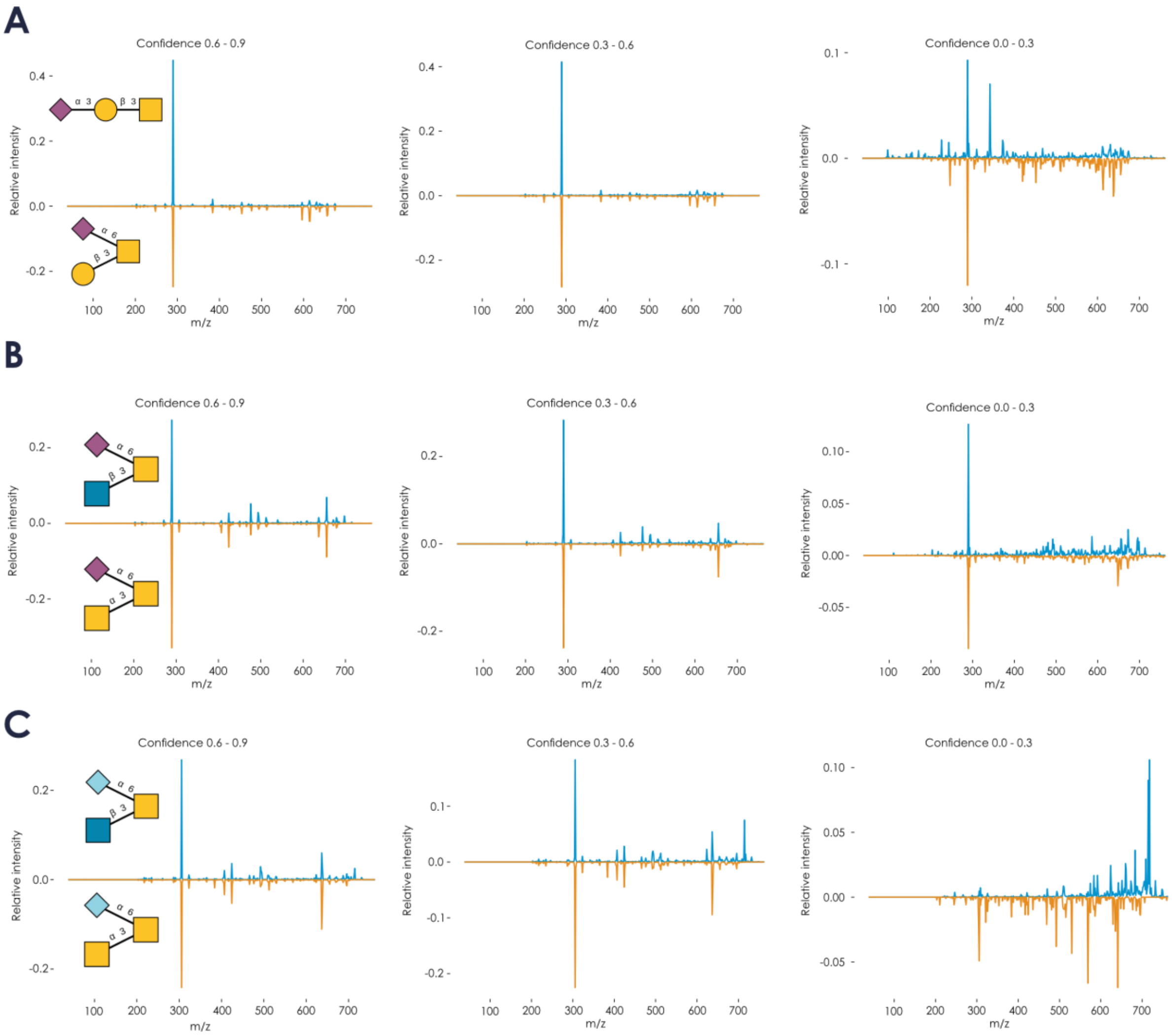


**Supplementary Figure 10. Loss of distinctive fragmentation with decreasing spectrum quality. A-C)** Similar to Figure 2, averaged spectra of Neu5Acα2-3Galβ1-3GalNAc / Galβ1-3(Neu5Acα2-6)GalNAc (A), GlcNAcβ1-3(Neu5Acα2-6)GalNAc / GalNAcα1-3(Neu5Acα2-6)GalNAc (B), and GlcNAcβ1-3(Neu5Gcα2-6)GalNAc / GalNAcα1-3(Neu5Gcα2-6)GalNAc (C) are juxtaposed for several bins of prediction confidence.

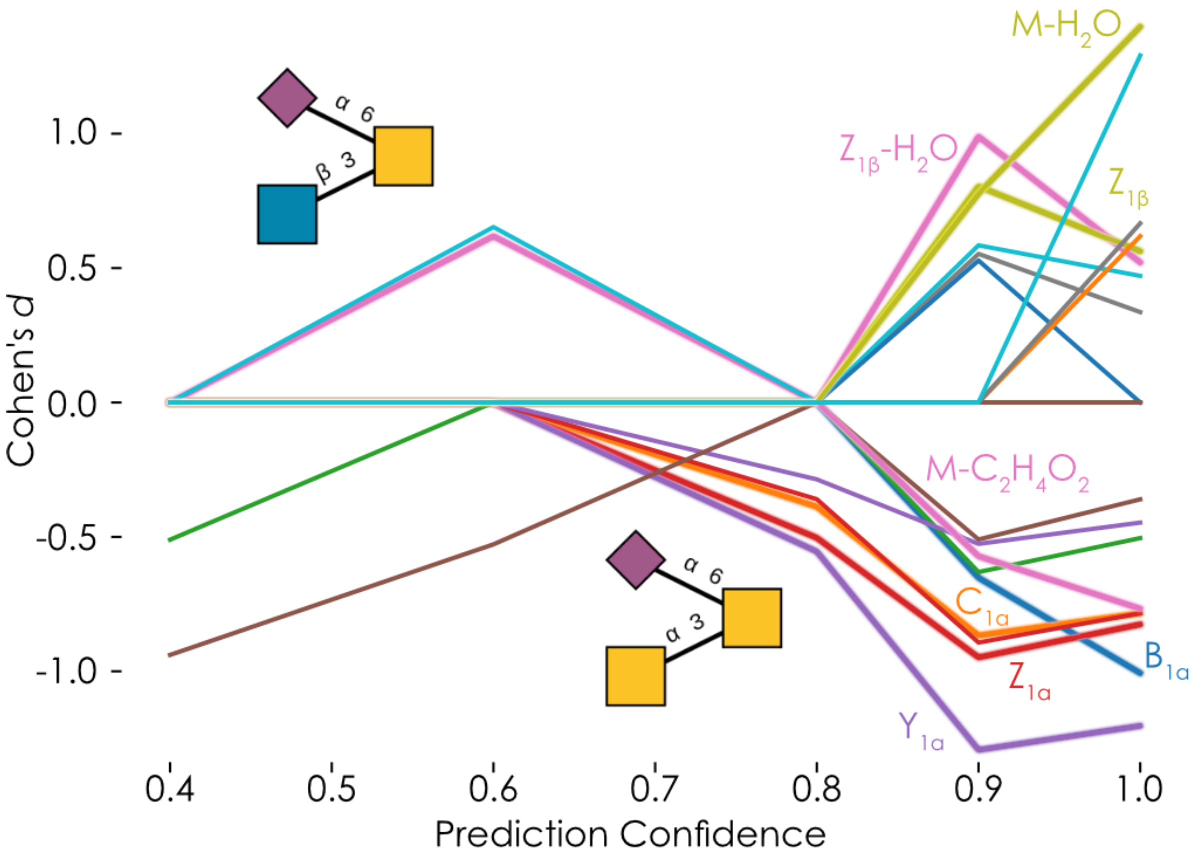


**Supplementary Figure 11. Signal strength of diagnostic ions across spectrum quality.** For the comparison of GlcNAcβ1-3(Neu5Acα2-6)GalNAc / GalNAcα1-3(Neu5Acα2-6)GalNAc, we binned spectra according to prediction confidence, as a measure of spectrum quality, and calculated the effect sizes of significantly different peaks in each bin. Significance was established via Welch's t-test of the normalized peak intensities ($p < 0.05$), with a Holm-Šídák correction for multiple testing. Effect size was calculated as Cohen's *d*. Positive effect size means more prevalent in the first structure. Line colors distinguish the different fragments. Example fragments, annotated via CandyCrumbs, are noted next to their line plots in Domon-Costello nomenclature.

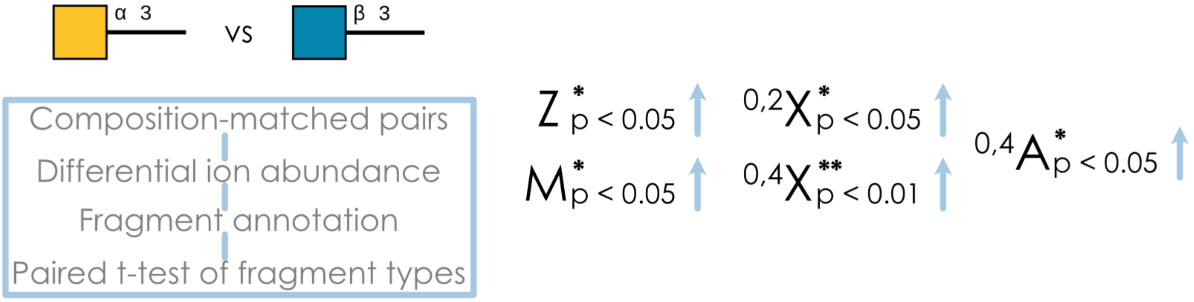


**Supplementary Figure 12. Systematic fragmentation differences in GalNAcα1-3 or GlcNAcβ1-3 containing *O*-glycans.** All *O*-glycan spectra containing GalNAcα1-3 or GlcNAcβ1-3 with confidence between 0.7 and 1.0 were used in the described workflow to ascertain their characteristic types of fragmentation.

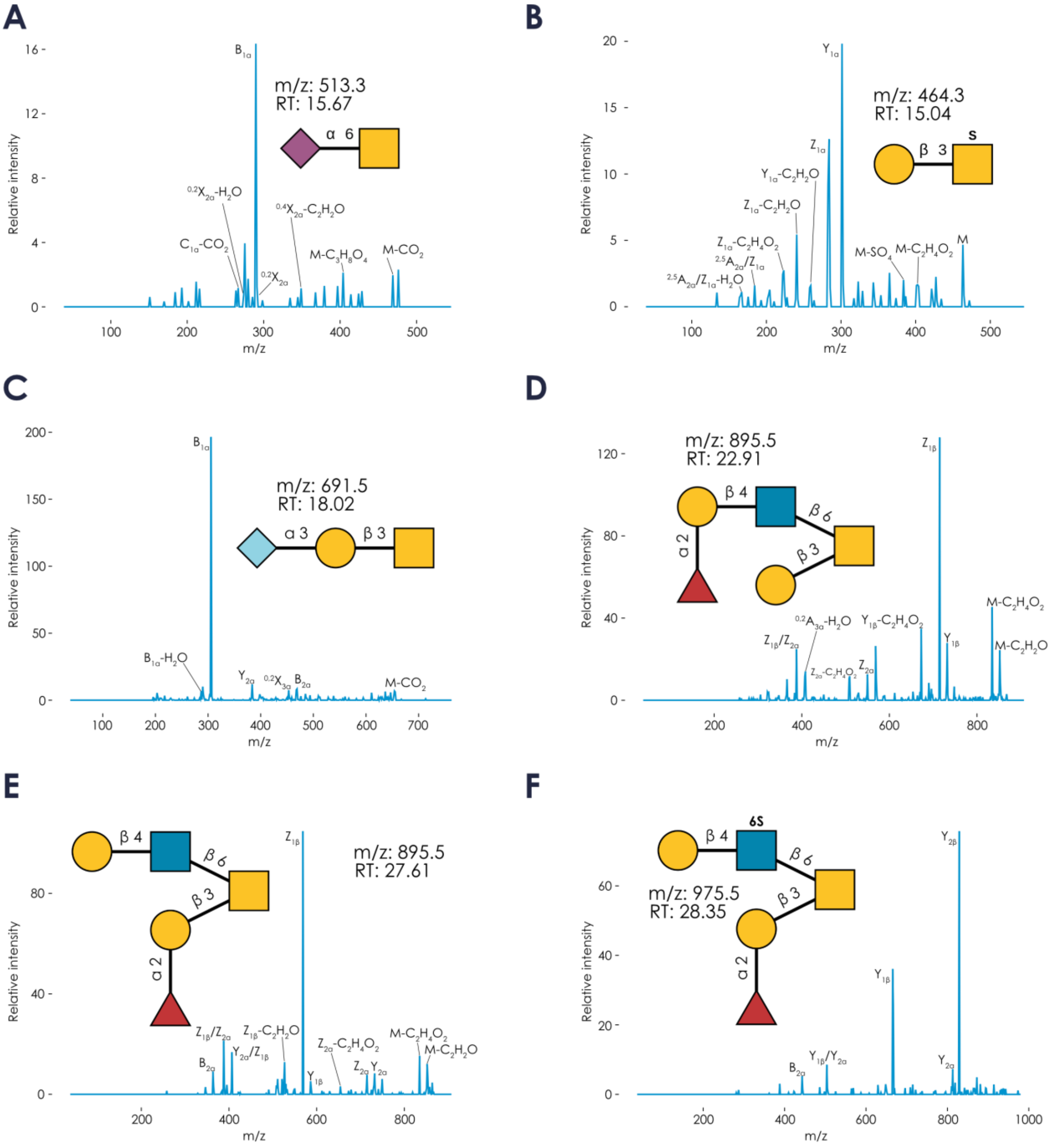

**Supplementary Figure 13. Increasing annotation comprehensiveness via CandyCrunch and CandyCrumbs. A-F)** We used CandyCrunch to predict all glycan structures from the file TT_200116Hayden_5 in GPST000307, containing murine intestinal *O*-glycans. We then chose example predictions which were absent from the associated annotation on GPST000307. The associated MS/MS spectra used for prediction, together with their m/z values, retention times, and predicted structures are shown with their fragments, annotated by CandyCrumbs, in the Domon-Costello nomenclature.

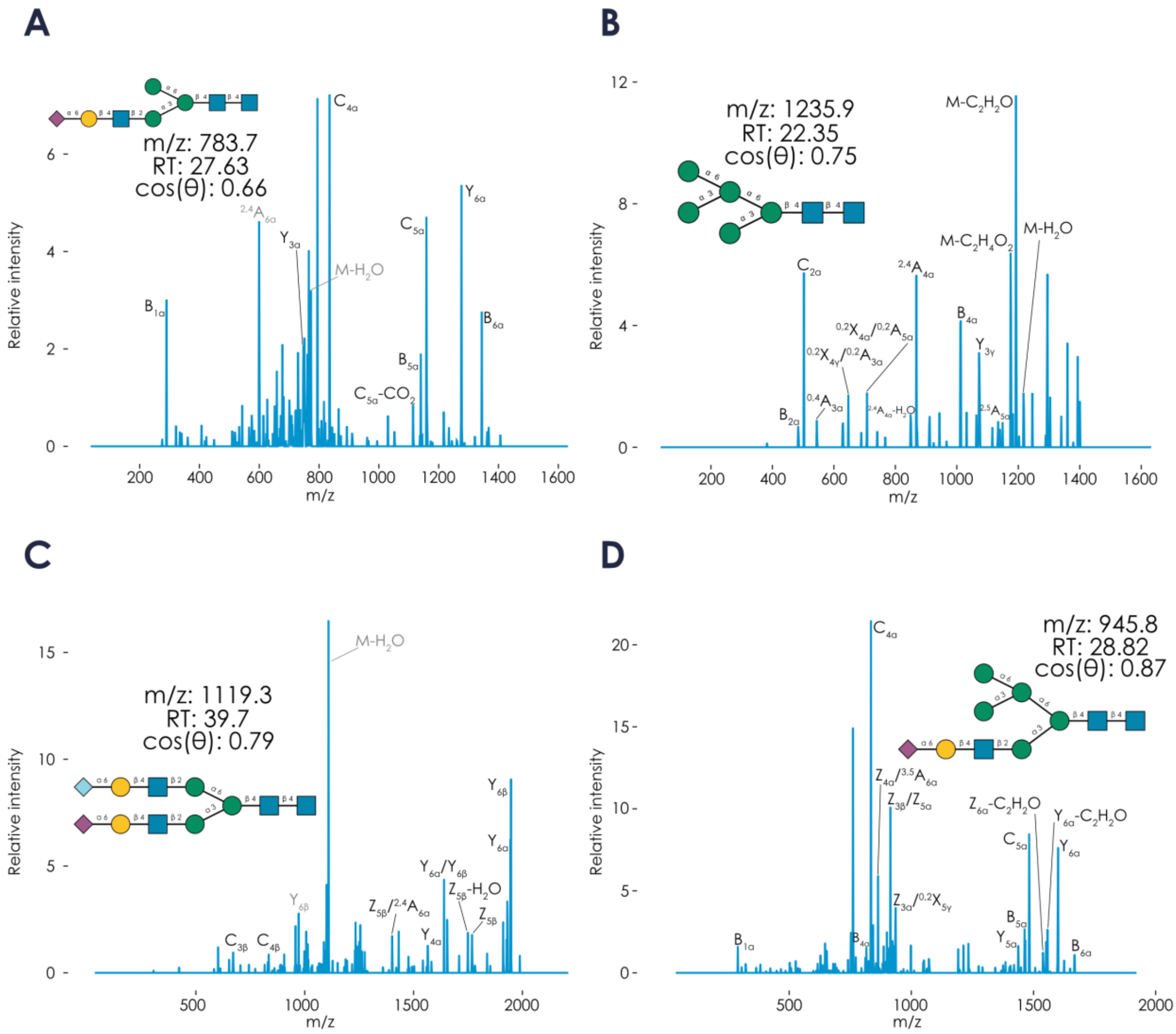


**Supplementary Figure 14. Predicting *N*-glycan structures from bluefin tuna. A-D)** Using the raw file LC_SBT_blood_NG_TR1_22082019_BB3_01_2514.d.mzXML from GPST000182, measuring reduced *N*-glycans from southern bluefin tuna (*Thunnus maccoyii*) blood in negative ion mode, we used our CandyCrunch pipeline to predict and curate *N*-glycan structures. Example predictions are shown with their m/z value and retention time in minutes, together with their $MS^2$ spectrum, with abundant fragments annotated in Domon-Costello nomenclature. Doubly-charged fragment ions are denoted with gray text. Also shown is the cosine similarity, cos(θ), of the shown spectrum and the averaged spectrum of all negative ion mode spectra of reduced glycans of the predicted structure with a confidence above 0.5 (see Fig. 2 for background).

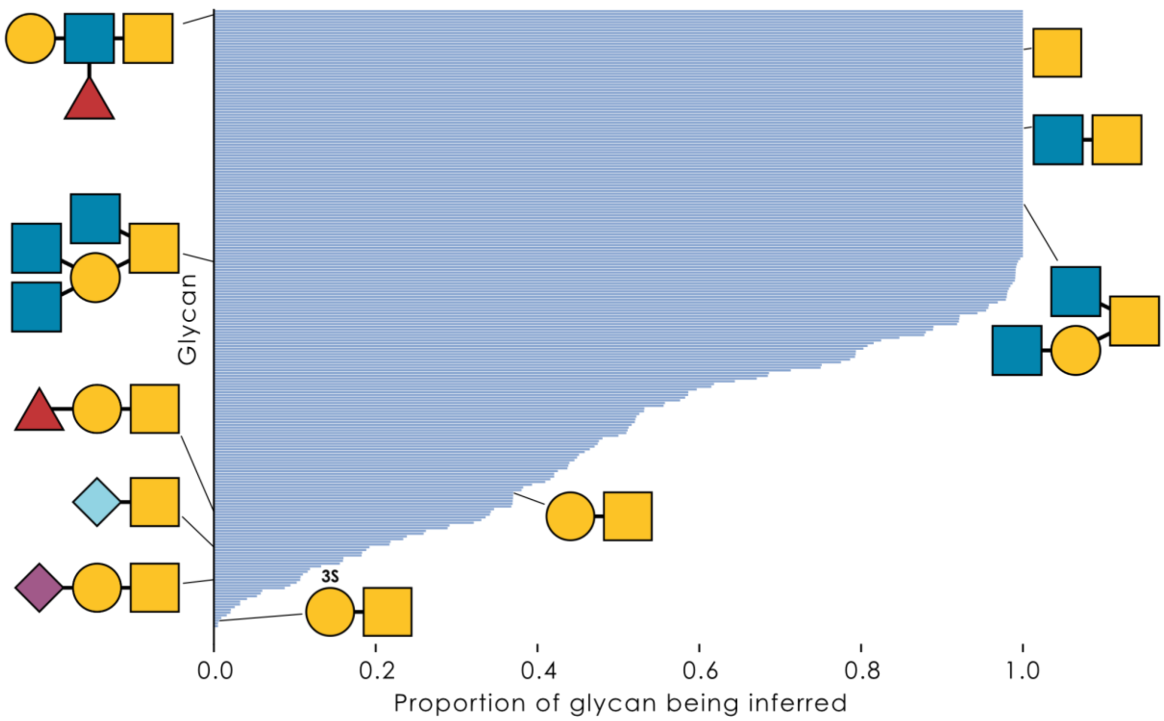


**Supplementary Figure 15. Analysis of inferred intermediates in biosynthetic networks of *O*-glycomes.** For all 1,003 biosynthetic networks we calculated from our *O*-glycome samples, we quantified whether a glycan was being present as an observed structure or an inferred intermediate to complete the network. We then calculated the ratio of these two quantities as an indicator of how likely it was to observe a given glycan via LC-MS/MS. Shown are the glycans that were present in the networks of at least 100 samples in our dataset, with example structures annotated via the SNFG.

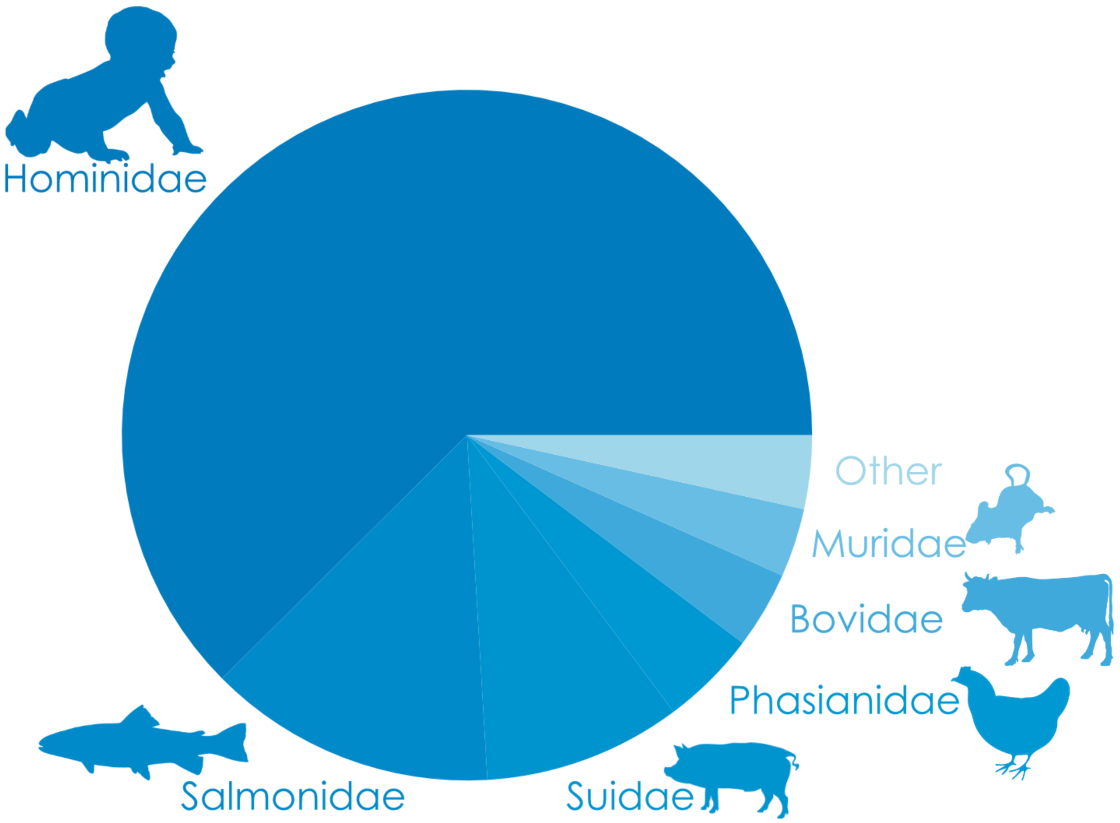


**Supplementary Figure 16. Taxonomic distribution of our dataset.** For our entire dataset of ~500,000 annotated mass spectra, we labeled each spectrum with the species it stemmed from and catalogued this information at the taxonomic family level, displaying relative proportions for the entire dataset. Taxonomic groups that are only distantly related to the displayed groups are likely to yield poorer predictions, due to stronger deviations from glycans seen during training.
